# Identification and Overexpression of Endogenous Transcription Factors to Enhance Lipid Accumulation in the Commercially Relevant Species *Chlamydomonas pacifica*

**DOI:** 10.1101/2025.05.01.651737

**Authors:** Abhishek Gupta, Aaron Oliver, João Vitor Dutra Molino, Kathryn MJ Wnuk-Fink, Marissa Tessman, Kalisa Kang, Évellin do Espirito Santo, Yasin Torres-Tiji, Michael D. Burkart, Stephen Mayfield

## Abstract

Sustainable low-carbon energy solutions are critical to mitigating global carbon emissions. Algae-based platforms offer potential by converting carbon dioxide into valuable products while aiding carbon sequestration. However, scaling algae cultivation faces challenges like contamination in outdoor systems. Previously, our lab evolved *Chlamydomonas pacifica*, an extremophile green alga, which tolerates high temperature, pH, salinity, and light, making it ideal for large-scale bioproduct production, including biodiesel. Here, we enhanced lipid accumulation in evolved *C. pacifica* by identifying and overexpressing key endogenous transcription factors through genome-wide in-silico analysis and in-vivo testing. These factors include Lipid Remodeling Regulator 1 (CpaLRL1), Nitrogen Response Regulator 1 (CpaNRR1), Compromised Hydrolysis of Triacylglycerols 7 (CpaCHT7), and Phosphorus Starvation Response 1 (CpaPSR1). Under nitrogen deprivation, CpaLRL1, CpaNRR1, and CpaCHT7 overexpression enhanced lipid accumulation compared to wildtype. However, CpaPSR1 increased lipid accumulation compared to wildtype in normal media despite causing no effect under nitrogen depravation, highlighting the difference in function based on media conditions. Notably, lipid analysis of CpaPSR1 under normal media conditions revealed a 2.4-fold increase in triglycerides (TAGs) compared to the wild type, highlighting its potential for biodiesel production. This approach provides a framework for transcription factor-focused metabolic engineering in algae, advancing bioenergy and biomaterial production.

**Graphical Abstract:** 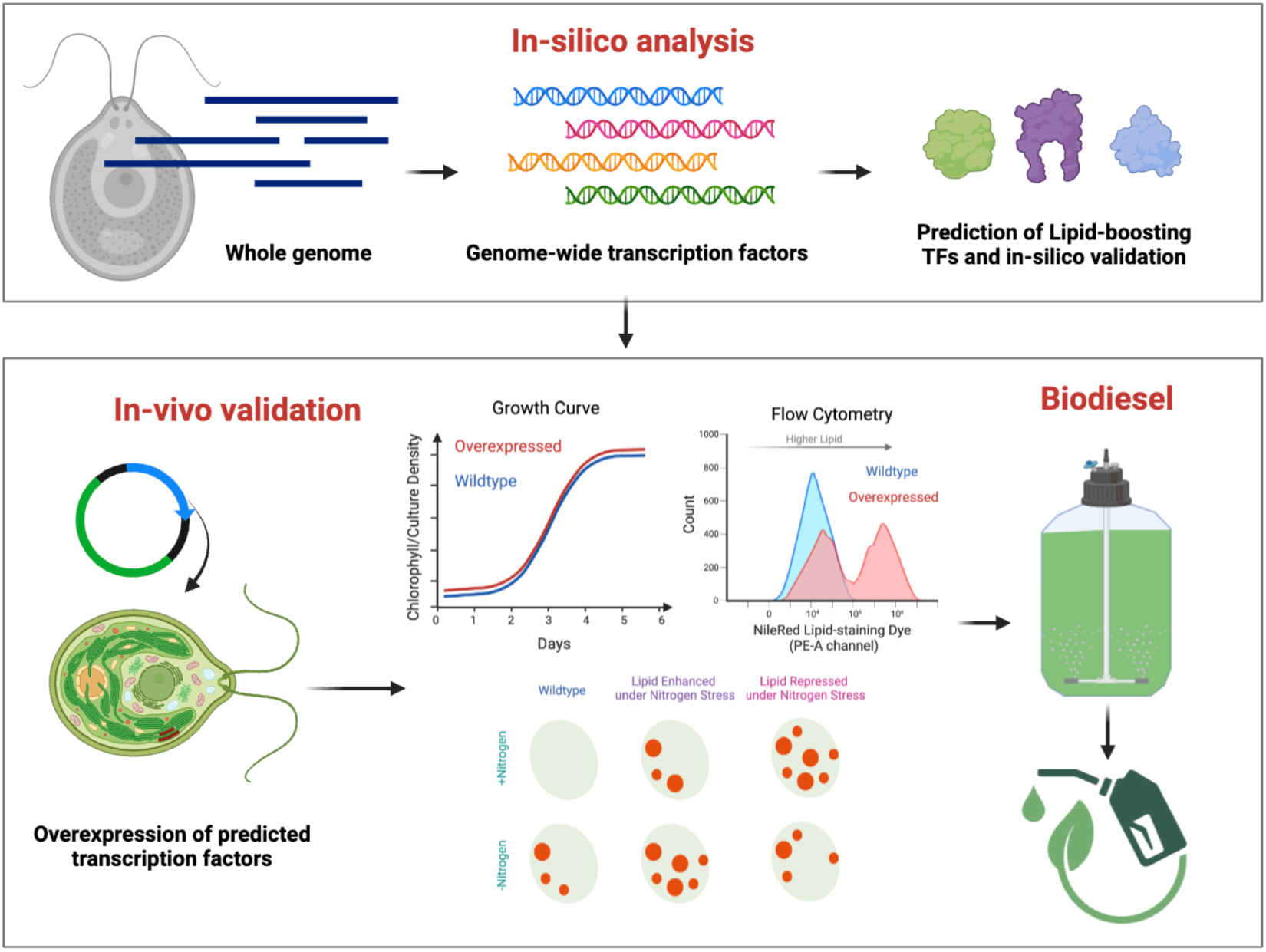

Created with BioRender.com.

## Introduction

Since the dawn of industrialization, the extensive utilization of fossil fuels has been a cornerstone of economic growth and technological advancement^1^. However, this heavy reliance has come at a significant cost to the environment, contributing to the current threat of climate change brought about by the emissions of greenhouse gases^2^. The resultant global warming, characterized by rising temperatures, melting ice caps, and extreme weather phenomena, underscores a pressing need to transition toward sustainable energy sources^3^. The shift towards sustainable energy is not merely a response to the environmental crisis but also a prudent approach to preparing for a future where fossil fuel resources are bound to become scarcer and more expensive^4^.

Microalgae hold enormous potential as a sustainable source of food, feed, and energy, presenting several advantages over traditional plant-based sources^5,6^. Their biomass is nutritionally rich, containing essential components such as lipids, proteins, and carbohydrates crucial for both human and animal nutrition. Some strains can accumulate up to 65-70% protein, making them an excellent source of nutritional protein, while others are rich in lipids, providing a sustainable source of omega-3 fatty acids^7^. Microalgae exhibit faster growth rates than conventional plants and can be harvested throughout the year without the need to compete for arable land^8^. Moreover, microalgae are extremely efficient in absorbing carbon dioxide and converting it to useful bioproducts much faster than most traditional crop plants^9^. These traits make microalgae an attractive feedstock for producing biodiesel and Sustainable Aviation Fuel^10^. In spite of this potential, the economic viability of microalgal products is challenged by the substantial costs tied to biomass production, encompassing expensive cultivation systems, harvesting, and processing, all of which inhibit its commercial practicality^11,12^.

To minimize the costs associated with biomass production and promote the adoption of microalgae-based technologies, it is essential to boost productivity both at the cellular and cultivation scales^13,14^. Increasing cellular productivity necessitates strain improvement, either through molecular modifications or breeding and selection, while rising biomass productivity on large-scale cultivation systems can be achieved through the use of strains with enhanced productivity as well as resilience to pond crashes^6,15,16^. Transcription factors (TFs) have been demonstrated to play a pivotal role in enhancing productivity in terrestrial crops like wheat and corn^17,18^. Transcription factors, proteins that oversee gene expression, can be leveraged to steer metabolic flux in a desired direction, thereby elevating biomass quality. Utilizing transcription factor-based metabolic engineering has been shown to significantly improve the quality of biomass production in microalgae^19^. For instance, several studies have employed native or plant transcription factors to augment lipids or starch production in various microalgae strains^20–23^. Extremophile strains have evolved to endure harsh conditions and have an advantage in reduced contamination risks due to limited resource competition and predator competition in extreme environments^24,25^. Hence, implementing transcription-factor-based metabolic engineering on an extremophile alga could be a robust approach to make microalgae a viable platform for advancing a sustainable economy^26–28^.

Recently, our group identified a unique green microalga, *Chlamydomonas pacifica*, capable of thriving in high-pH, high-temperature, and high-salinity conditions (CC-5697 & CC-5699)^29^. Moreover, we have also evolved this strain to have a high tolerance to light (CC-6190, 402wt x 403wt)^30^. The newly available genomic resources for this strain make it possible to conduct computational studies of transcription factors pertinent to genetic engineering. In this study, we enhanced lipid accumulation in the evolved *C. pacifica* by identifying and overexpressing endogenous transcription factors. Through a comprehensive genome-wide in-silico analysis and by in-vivo testing, we identified key endogenous putative transcription factors, including CpaLRL1, CpaNRR1, CpaCHT7, and CpaPSR1 in *C. pacifica*. Overexpression of these transcription factors induced lipid accumulation under normal minimal or nitrogen-deplete media for all four transgenic lines. Additionally, we converted the extracted lipids into biodiesel, underscoring the commercial relevance of these transgenic high-lipid strains. This approach not only positions these strains as prime candidates for large-scale biodiesel and biomaterial production but also provides a framework for applying transcription factor-focused metabolic engineering to other commercially important microalgae species.

## Results

### Overview and comparative analysis of transcription factors in *C. pacifica* with other algae, and higher plants

The identification and classification of transcription factors depend significantly upon their unique features. Essential attributes, like DNA-binding domains and specific non-binding areas, are crucial for recognizing TFs^31,32^. These elements are fundamental in DNA interaction and gene regulation, which are the primary functions of TFs. Additionally, TFs are grouped into various families based on these characteristics, with each family having distinct domains that aid in distinguishing between them and shedding light on their specific roles and regulatory processes. The transcription factor database PlantTFDB has established rules for family classification by extensively reviewing existing literature^33^. This database contains transcription factors for 165 species across 9 taxonomic groups encompassing plants and algae.

To predict all transcription factors, we employed the transcription factor prediction tool from PlantTFDB, leveraging its methodology to detect TFs in the *C. pacifica* genome. Given the abundance of chlorophyte genomes within the PlantTFDB database (16 in total), this resource is apt for forecasting TFs in the green alga *C. pacifica*. This initial method enabled us to successfully predict a total of 162 transcription factors in *C. pacifica* (**Supplementary Figure 1; Supplementary Table S1**). These potential transcription factors were additionally annotated using the Clusters of Orthologous Genes (COG) database to identify broad functional potential (**Figure 1a**). This analysis assigned 48% of transcription factors to a functional category based on this analysis. The most common COG categories for our transcription factors are associated with posttranslational modifications and signal transduction. Regulation of these pathways will define *C. pacifica* response to abiotic stresses. Few transcription factors associated with lipid and starch metabolism could be identified with this initial analysis, requiring additional techniques to predict which transcription factors would be most relevant for exploiting *C. pacifica* as a source of food and fuel.

**Figure 1:**
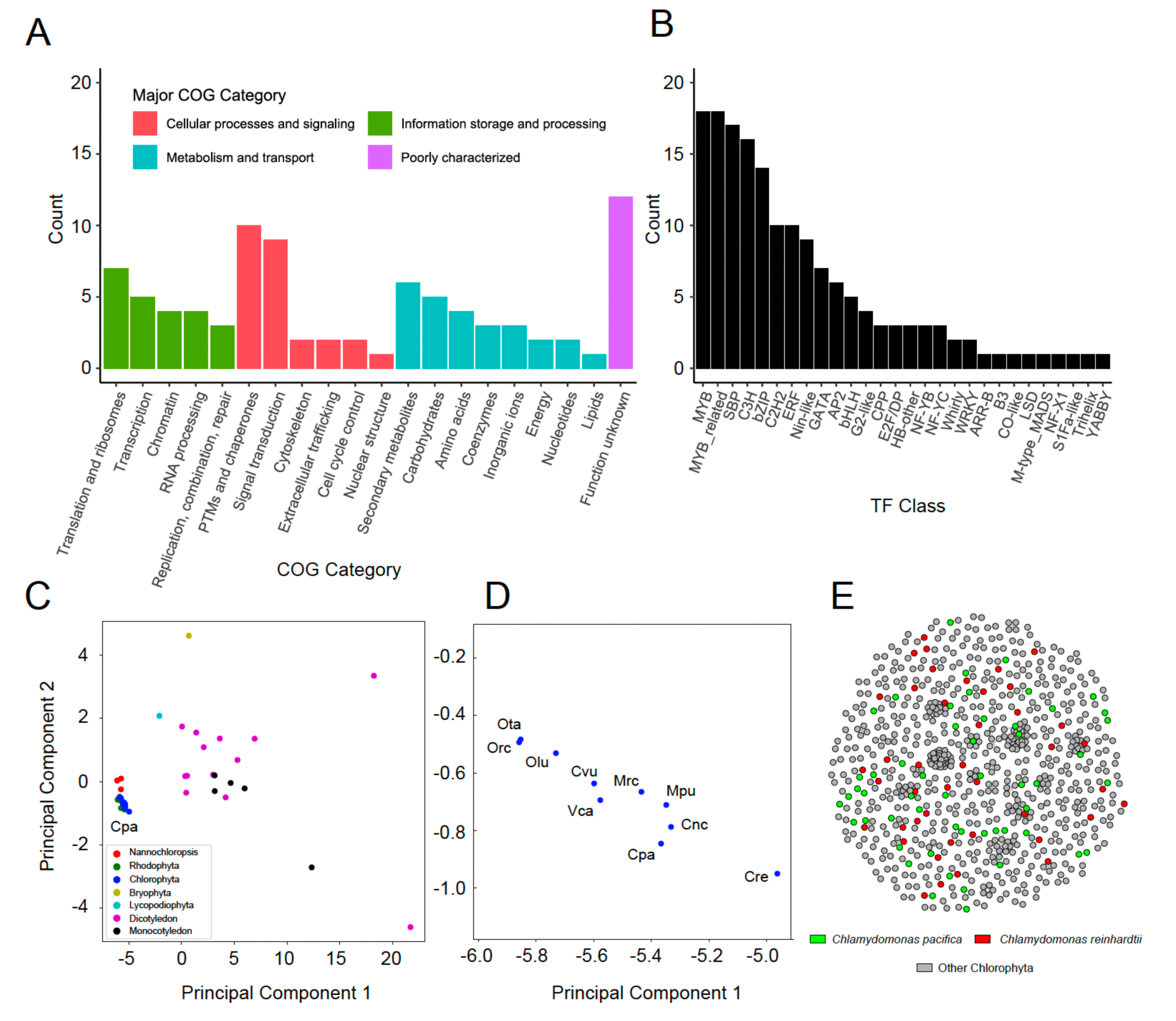
Overview and comparative analysis of transcription factors in *Chlamydomonas pacifica* with other algae and higher plant species. (**A**) Distribution of transcription factors across different Clusters of Orthologous Genes (COG) categories, indicating their potential functions in *Chlamydomonas pacifica*. (**B**) Bar chart showing the distribution of transcription factors across various families in *Chlamydomonas pacifica*, highlighting the abundance of different types within the species. (**C**) PCA plot illustrating the transcription factor profiles of *Chlamydomonas pacifica* compared with those of various other algae and higher plant species. (**D**) A zoomed-in PCA plot focusing on the Chlorophyta group, highlighting the close relationship of *Chlamydomonas pacifica* with other green algae species. (**E**) Network plot showing the interactions between transcription factors of *Chlamydomonas pacifica*, *Chlamydomonas reinhardtii*, and other Chlorophyta species. The labels for species in panel (**C,D**) are as follows: *Chlamydomonas reinhardtii* (Cre), *Chlamydomonas pacifica* (Cpa), *Volvox carteri* (Vca), *Chlorella sp. NC64A* (Cnc), *Coccomyxa sp. C-169* (Cvu), *Micromonas pusilla CCMP1545* (Mpu), *Micromonas sp. RCC299* (Mrc), *Ostreococcus lucimarinus CCE9901* (Olu), *Ostreococcus sp. RCC809* (Orc), and *Ostreococcus tauri* (Ota).

Our findings particularly highlight the prevalence of certain TF families—namely Myeloblastosis (MYB; 11%), MYB-related (11%), SQUAMOSA promoter binding protein (SBP; ∼10%), Cysteine3Histidine (C3H; ∼10%), and Basic leucine zipper (bZIP; ∼9%)—which together represent over half (51.23%) of all identified transcription factors (**Figure 1b**). MYB, SBP, and bZIP TFs have been identified as key players in lipid metabolism and stress responses, instrumental in enhancing microalgae’s potential for biofuel production^22,34–39^. MYB, MYB-related, and C3H transcription factors are known to be involved in the transcription regulation of astaxanthin synthesis-related genes in microalgae *Haematococcus pluvialis*^40^. Moreover, C3H TFs are known to be involved in biotic and abiotic stress^41,42^.

Grasping the evolutionary and functional landscapes of transcription factors within microalgae, and their comparison with other algae species and higher plant species, is fundamental to deciphering their relational and adaptive intricacies. Here, we performed Principal Component Analysis (PCA) to compare the transcription factor profiles of *C. pacifica*, a newly identified green algae, with those of other algae such as *Nannochloropsis* and Rhodophyta (red algae), as well as a range of higher plant species, encompassing a total of 36 species and 58 transcription factor families. PCA results demonstrated that *C. pacifica* clusters closely with other Chlorophyta (green algae) species (**Figure 1c**). A statistical t-test to assess the segregation between algae and higher plant species revealing a significant difference between these two groups (PC1: t-test statistic: -7.04, p-value: ∼3.37e-08; PC2: t-test statistic: -2.24, p-value: ∼0.032). These findings suggest a distinct segregation between the algae group and the higher plant species in our PCA analysis. An analysis of Chlorophyta highlighted *C. pacifica’s* transcription factor profile as closely related to other green algae, particularly *C. reinhardtii* (**Figure 1d**). Spearman’s rank correlation (r = 0.947, p-value = 3e-29) confirmed their strong TF profile similarity, suggesting both a shared evolutionary history as well as shared regulatory mechanisms.

Subsequently, a network plot was constructed to examine the clustering patterns of *C. pacifica* transcription factors alongside those from other Chlorophyte genomes, such as *C. reinhardtii* (**Figure 1e**). Of the 162 *C. pacifica* transcription factors predicted by PlantTFDB, just 17% of them were found in a cluster containing a transcription factor from *C. reinhardtii*. An additional Sankey plot was constructed to elucidate the transcription factor profile similarities and differences between *C. pacifica* and *C. reinhardtii* (**Supplementary Figure 2**). Notably, the Sankey plot revealed the absence of predicted Heat Shock Factors (HSF) and DNA binding with one finger (*Dof*) TF families in *C. pacifica*, which are present in *C. reinhardtii*. The observed discrepancies could arise from limitations in the prediction technique, inaccuracies in genome assembly or sequencing, or the actual absence of these transcription factor families, underscoring the need for further investigation.

### High-throughput approach to identity lipid enhancing transcription factors

To pinpoint TFs with the potential to boost lipid production, we synthesized data from multiple transcription factor prediction tools (**Figure 2**, see **Methods**). We started by identifying all TFs using two additional databases, iTAK and TAPscan, alongside PlantTFDB. iTAK and TAPscan identified 182 and 183 TFs, respectively **(Supplementary Tables S2-S3**). Subsequently, we selected only those TFs consistently predicted by all three databases, yielding 139 common TFs (**Supplementary Table S4**). Next, these commonly predicted TFs were compared via phylogenetic trees and active domain alignment with a list of known TFs involved in lipid metabolism, as reported in the existing literature^21^. This identified 10 transcription factors in *C. pacifica* that are orthologs of TFs found in *Arabidopsis thaliana*, *Auxenochlorella protothecoides*, *C. reinhardtii*, *Dunaliella parva*, and *Scenedesmus obliquus* (**Figure 2**, **Table 1**). These TFs are categorized into the AP2, NF-YB, MYB, bZIP, CPP, and SBP families.

**Figure 2:**
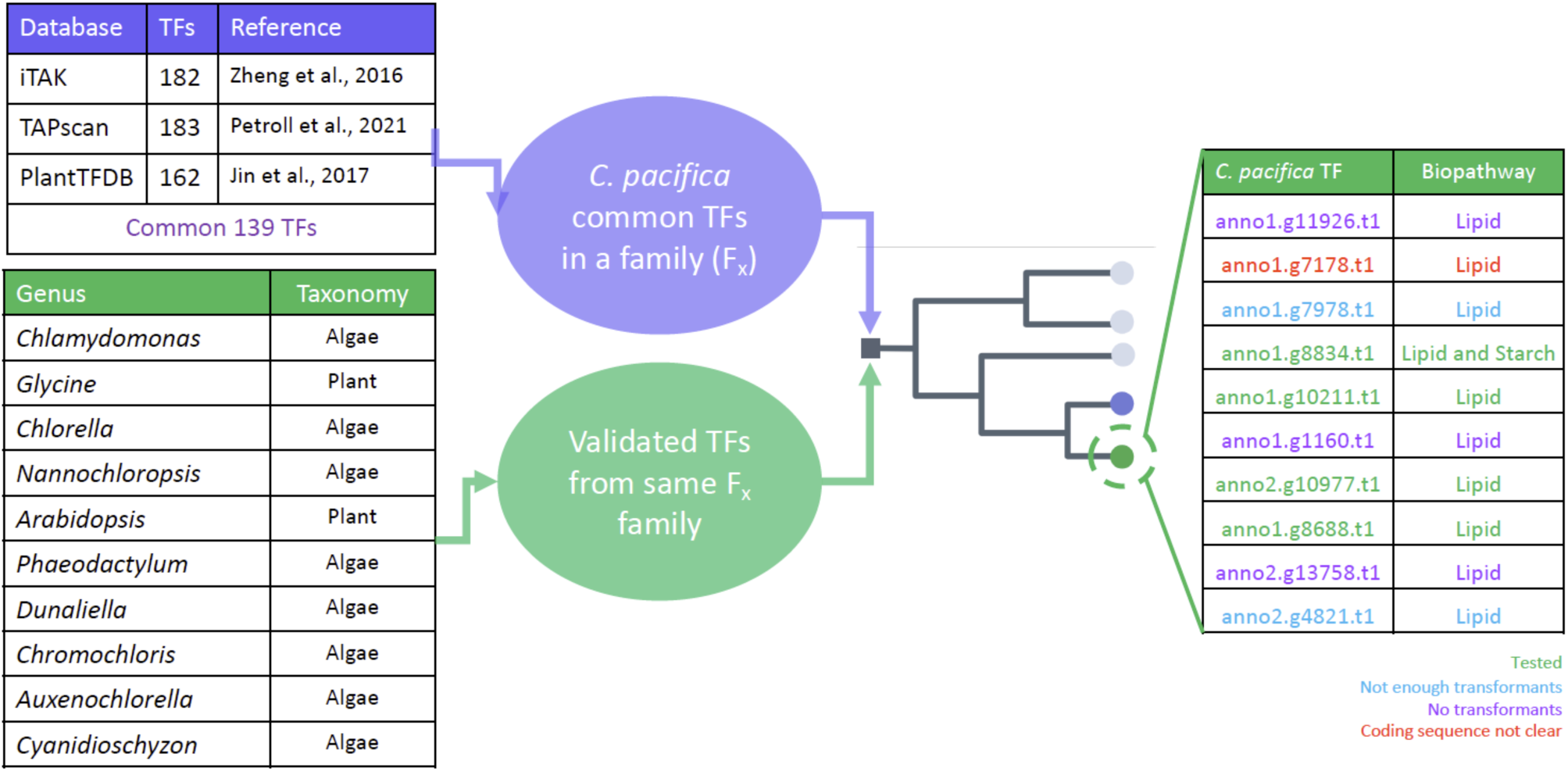
Pipeline for predicting transcription factors involved in starch and lipid biosynthesis in *C. pacifica*. The schematic illustrates the prediction of transcription factors involved in starch and lipid biosynthesis in *C. pacifica*. In Step 1, transcription factors were predicted using three different tools, and a comprehensive database of validated transcription factors was created from 10 different genera based on literature. In Step 2, a phylogenetic tree analysis was conducted on the transcription factors from *C. pacifica* and the curated validated database within the same transcription factor family. This analysis resulted in the identification of 10 predicted transcription factors involved in the biosynthetic pathways. The color coding of various transcription factors signifies whether they have been experimentally tested in this study.

**Table 1:** Predicted transcription factors associated with lipid and starch synthesis enhancement in *Chlamydomonas pacifica*.

| Microalgae Species | Source Species | Transcription Factor | Family | *C. pacifica* Candidate | Biopathway | Reference |
| --- | --- | --- | --- | --- | --- | --- |
| *Nannochloropsis salina* | *Arabidopsis thaliana* | WRINKLED1 (AtWRI1) | AP2 | anno1.g11926.t1 | Lipid | Kang et al., 2017 |
| *Chlorella ellipsoidea* | *Arabidopsis thaliana* | Leafy Cotyledon1 (AtLEC1) | NF-YB | anno1.g7178.t1 | Lipid | Liu et al., 2021 |
| *Auxenochlorella protothecoides* | *Auxenochlorella protothecoides* | Myeloblastosis 6 (ApMYB6) | MYB | anno1.g7978.t1 | Lipid | Xing et al., 2021 |
| *Chlamydomonas reinhardtii* | *Chlamydomonas reinhardtii* | Phosphorus Starvation Response 1 (PSR1) | MYB | anno1.g8834.t1 | Lipid and Starch | Bajhaiya et al., 2016;<br>Ngan et al., 2015 |
| *Chlamydomonas reinhardtii* | *Chlamydomonas reinhardtii* | Lipid Remodeling Regulator 1 (LRL1) | MYB | anno1.g10211.t1 | Lipid | Hidayati et al., 2019 |
| *Chlamydomonas reinhardtii* | *Chlamydomonas reinhardtii* | Basic Leucine Zipper 1 (CrbZIP1) | bZIP | anno1.g1160.t1 | Lipid | Yamaoka et al., 2019 |
| *Chlamydomonas reinhardtii* | *Chlamydomonas reinhardtii* | Compromised Hydrolysis of Triacylglycerols 7 (CHT7) | CPP | anno2.g10977.t1 | Lipid | Tsai 2014 |
| *Chlamydomonas reinhardtii* | *Chlamydomonas reinhardtii* | Nitrogen Response | SBP | anno1.g8688.t1 | Lipid | Boyle et al., 2012 |
|  |  | Regulator 1<br>(NRR1) |  |  |  |  |
| *Dunaliella parva* | *Dunaliella parva* | WRINKLED1<br>(DpWRI1) | AP2 | anno2.g13758.t1 | Lipid | <u>Shang et al., 2016</u> |
| *Scenedesmus obliquus* | *Scenedesmus obliquus* | SQUAMOSA-promoter binding<br>protein-like 12<br>(SoSPL12) | SBP | anno2.g4821.t1 | Lipid | <u>Calhoun et al., 2023</u> |

To enhance confidence in our prediction, we investigated several of the predicted transcription factors in-depth. For instance, our analysis identified a transcription factor in *C. pacifica* with significant resemblance to *C. reinhardtii’s* PSR1 and *Arabidopsis thaliana*’s PHR1, characterized by MYB and coiled-coil domains (**Figure 3a**; percentage identity = ∼63%, p < 1e-05, permutation-test). PSR1 is known for its regulatory function in lipid and starch biosynthesis during nitrogen and phosphorus deficiency in *C. reinhardtii*, and PHR1 is implicated in triacylglycerol accumulation under phosphorus starvation in *Arabidopsis thaliana*^43–45^. Furthermore, we have pinpointed another TF that bears resemblance to the lipid-enhancing WRINKLED1 (WRI1) transcription factor, as observed in *A. thaliana* and *Helianthus annuus* (**Supplementary Figure 3**; percentage identity = ∼56%, p < 1e-05, permutation-test) ^46,47^. This TF maintains the key characteristics of the AP2 DNA binding domain, the VYL transcriptional activation motif, and two phosphorylation sites essential for protein stability^48^.

**Figure 3:**
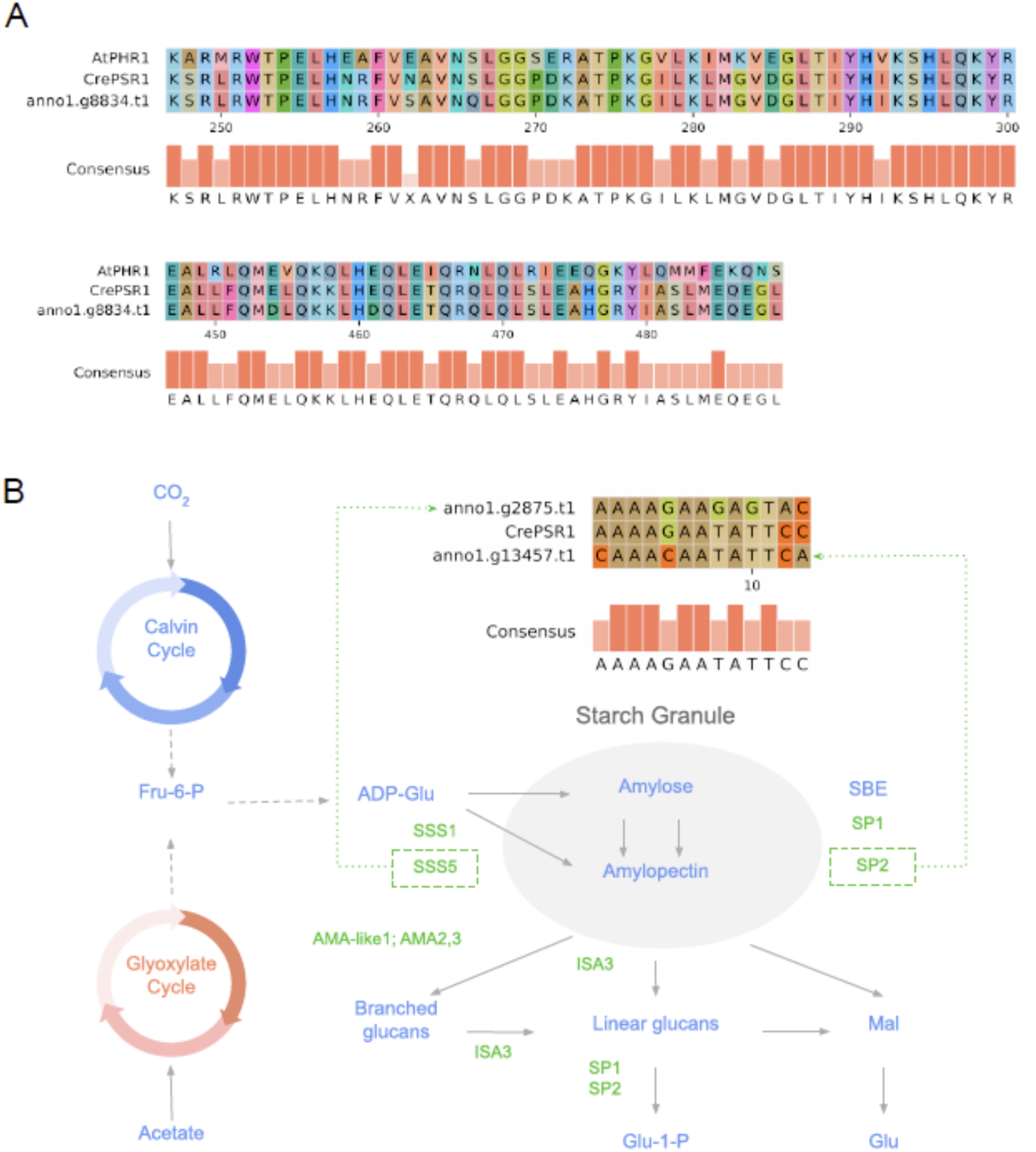
Conservation of transcription factor domain and genes in starch metabolism. (**A**) Alignment showing MYB and coiled-coil domains from *Chlamydomonas reinhardtii*’s PSR1 (CrePSR1), *Arabidopsis thaliana’s* PHR1 (AtPHR1), and an orthologous predicted CpaPSR1 transcription factor from *Chlamydomonas pacifica* (anno1.g8834.t1), highlighting amino acid conservation. (**B**) An illustration of orthologous genes related to starch synthesis and breakdown is marked in green text. It features the enriched motifs within the promoter sequences of two of these genes that are similar to the consensus motif sequence of CrePSR1. The schematic is adapted from Bajhaiya et al., 2016. The enzyme label SBE represents Starch Branching Enzyme, and the orthologous gene definitions are described in **Supplementary Table S5**.

As the *C. reinhardtii* PSR1 TF has been shown to affect is known starch metabolism, we extended our investigation to include orthologs of genes involved in the *C. reinhardtii* starch pathway^43^. In this pursuit, we successfully identified orthologs for all eight genes reported in the study by Bajhaiya et al. within the genome of *C. pacifica* (**Figure 3b, Supplementary Table S5**), whose functions were confirmed with EggNOG^43^. A noteworthy discovery emerged from examining the native promoters of these genes in *C. pacifica*. Two of these gene promoters exhibited motifs are strikingly similar to the consensus motif of PSR1 as identified in the PlantTFDB database (**Figure 3b**). This similarity in promoter Transcription factor binding sites (TFBSs) suggests a potential regulatory relationship with PSR1 in *C. pacifica* akin to that observed in *C. reinhardtii*.

### Overexpression of predicted endogenous transcription factors

To validate our predictions, we experimentally tested them in the evolved *C. pacifica* strain to determine if overexpression could enhance lipid accumulation. We cloned 9 out of 10 transcription factors under the β2 tubulin promoter sequence with hygromycin selection. One transcription factor gene (anno1.g7178.t1) was not well annotated and thus was not cloned (**Figure 2**). Among the 9 TFs, we successfully obtained transformants for 6, and only 4 provided enough transformants for screening. The absence of transformants in some TF lines may be attributed to using the upstream β2 tubulin promoter, whose strong and constitutive expression may prove to be lethal. To address this issue, employing an inducible promoter for the overexpression of TFs could provide a more controlled and effective approach. We proceeded with these 4 TFs (anno1.g10211: CpaLRL1, anno1.g8688: CpaNRR1, anno2.g10977: CpaCHT7, anno1.g8834: CpaPSR1) for further evaluation. We first confirmed the transformants through colony PCR and sequencing (see **Methods**; **Supplementary Figure 5**). Subsequently, RT-qPCR analysis revealed that transcript levels of CpaLRL1, CpaNRR1, CpaCHT7, and CpaPSR1 were elevated by 3.13-fold, 3.41-fold, 6.41-fold, and 10.74-fold, respectively, in their corresponding overexpression lines compared to the wild type under normal growth conditions.

First, we compared the growth of the transgenic lines to the wildtype evolved strain, as overexpression of a transcription factor can impede the growth of the cells. However, we observed no significant differences in growth (p > 0.05 ANOVA for all pairwise differences), as measured by culture density and chlorophyll fluorescence, between the transgenic lines and wild-type cells (**Figure 4**). Next, we evaluated lipid accumulation in the transgenic lines using Nile Red lipid-binding dye and flow cytometry in the phycoerythrin (PE) channel under normal and nitrogen-deprived minimal media conditions (**Figure 5a, b**). A higher response in this channel corresponds to greater lipid content in the stained cells. We compared the population percentage above a threshold of 7.7×10⁴ RFU (Relative Fluorescence Units). This threshold value was selected to more effectively distinguish a specific subpopulation of cells (see **Methods**). Approximately 3.32% and 22.5% of the wild-type population exceeded this threshold under normal and nitrogen-deprived conditions, respectively (**Figure 5a, b**). All the transcription factors resulted in statistically significant (Wilcoxon ranked-sum, p < 0.05) increase of lipid accumulation in minimal media. In addition, CpaLRL1, CpaNRR1, CpaCHT7, and CpaPSR1 respectively displayed 1.3, 2.8, 7.0, and 11.9 times more observations above the nile red cutoff threshold compared to the wildtype (**Figure 5a**). Under nitrogen-deprived conditions, lipid accumulation also increased in CpaLRL1, CpaNRR1, and CpaCHT7 but showed no statistically significant increase for CpaPSR1 compared to wildtype, suggesting different roles for these transcription factors under nitrogen stress (**Figure 5b**). Confocal microscopy corroborated our flow cytometry results, with visual inspection showing more lipid-producing cells in the transgenic strains compared to the wild type under both media conditions (**Figure 6**).

**Figure 4:**
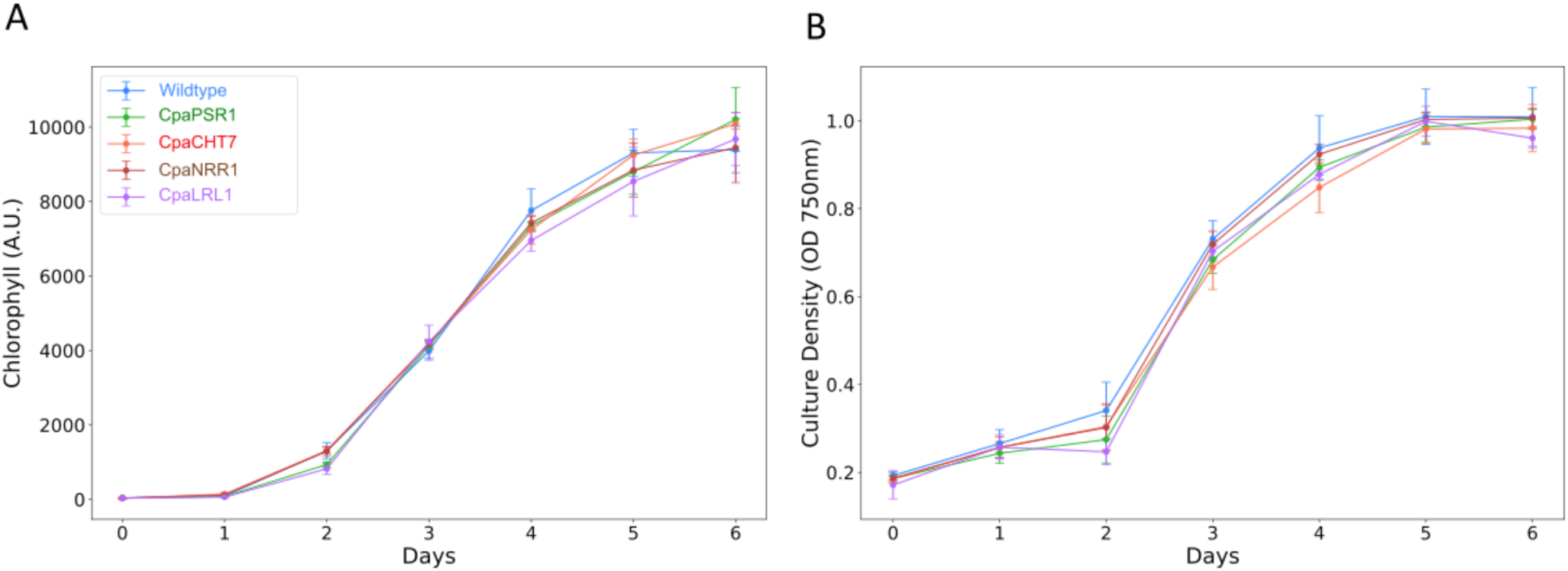
Cell growth measured as chlorophyll fluorescence and cell density in transgenic lines. (**A**), (**B**) Growth curves depicting chlorophyll levels and culture density of transcription factor overexpressed lines compared to the wildtype strain, respectively. Each line represents three replicates. A.U. stands for arbitrary units. O.D. stands for optical density.

**Figure 5:**
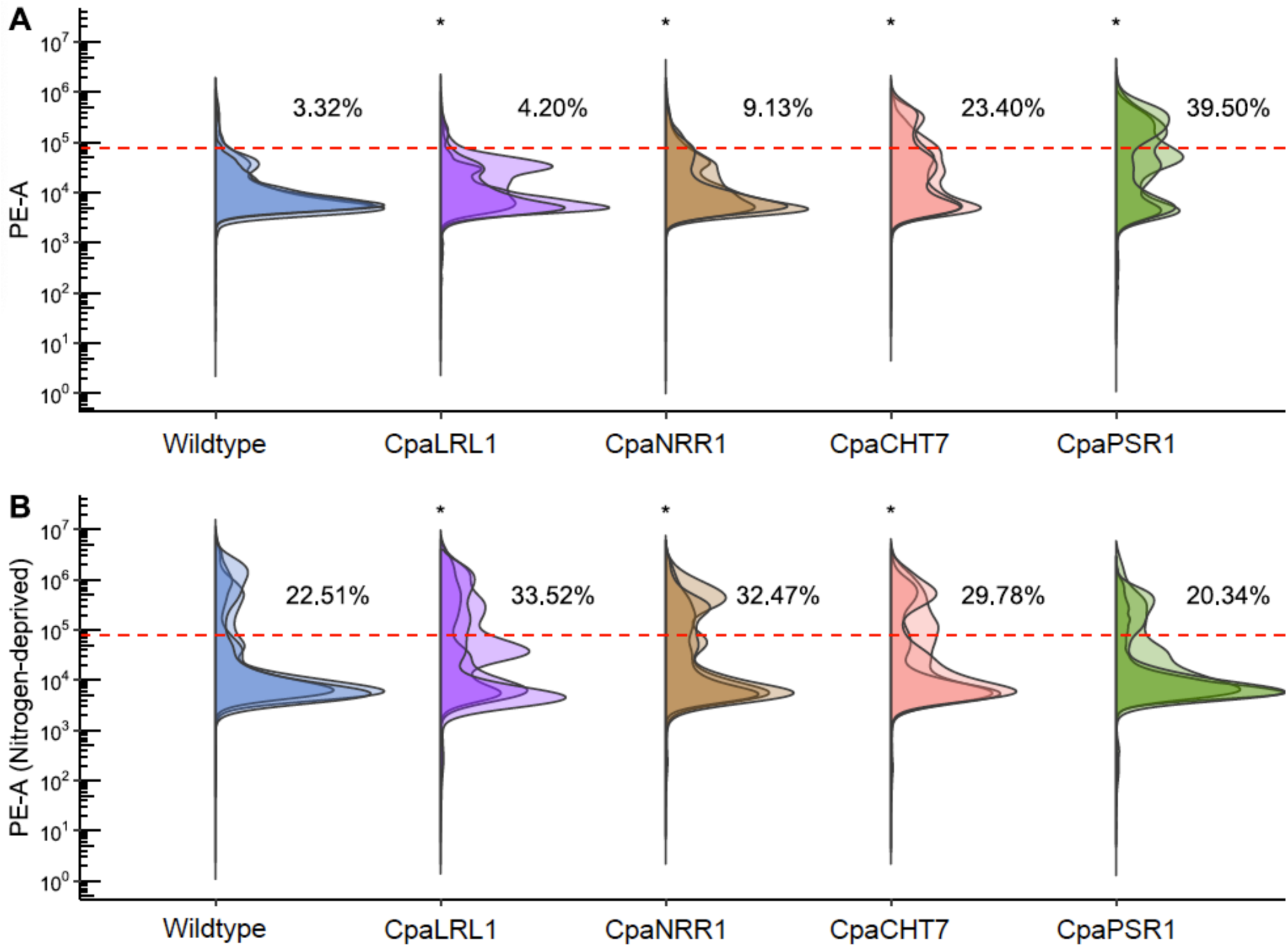
Flow cytometry data showing enhanced lipid production. (**A**), (**B**) Flow cytometry analysis of Nile red-stained cells was conducted to quantify lipid content in transcription factor overexpressed lines compared to the wildtype strain under normal minimal or nitrogen-deprived media conditions. The statistics displayed in the top right corner of each plot correspond to populations above the threshold of 7.7×10^4^. PE-A refers to the phycoerythrin-area channel. “Normal” and “-N” represent normal and nitrogen-deprived media conditions, respectively. Each plot shows curves corresponding to three biological replicates. ’*’ represents significant upward differences of the entire distribution from wildtype (Wilcoxon ranked sum test, p < 0.001) and each media is compared separately.

**Figure 6:**
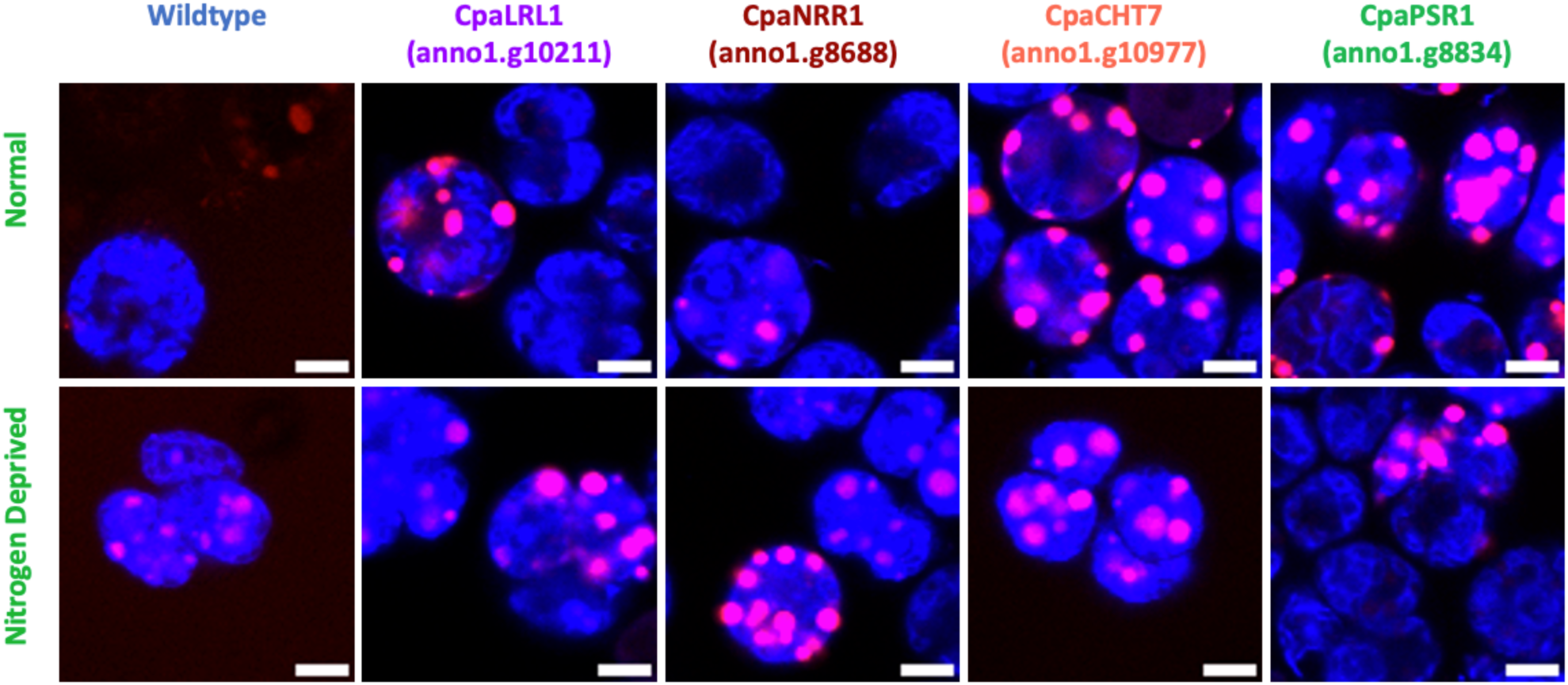
Confocal microscopy of cells in different media conditions. Cells in normal minimal and nitrogen-deprived media, respectively. The scale bar represents 5 µm. Blue and magenta correspond to Chlorophyll and triacylglycerol, respectively.

To further examine the differences in lipid accumulation between the wild-type and the promising CpaPSR1 under normal conditions, semi-quantitative lipidomic analysis was conducted on the two strains. Through this analysis, the relative abundance of triglycerides (TAGs) and diglycerides (DAGs) were determined for three replicates (see **Method**). For the wildtype, the total TAG content was found to be 152 ± 21 relative abundance/mg biomass, while the CpaPSR1 contained 369 ± 70 relative abundance/mg biomass. The diglyceride analysis showed values of 868 ± 128 relative abundance/mg biomass for the wildtype while the CpaPSR1 samples had DAG content of 241 ± 102 relative abundance/mg biomass. The observed increased accumulation of neutral TAGs in the CpaPSR1 sample has favorable implications for the cellular function and industrial relevance of this strain as TAGs provide more fatty acid chains per molecule compared to DAGs that can be used for biodiesel production.

Overall, the flow cytometry and microscopy, and lipidomics data validated our predictions and provided new insights into the functioning of these transcription factors under both normal and nitrogen-stress conditions. Notably, CpaPSR1 exhibited the highest lipid accumulation under normal media conditions, eliminating the need for time-consuming and resource-intensive nitrogen deprivation setups. This makes CpaPSR1 a prime candidate for large-scale biodiesel production.

### Biodiesel Production

To demonstrate commercial applications and understand potential alterations in fatty acids produced under varied growth conditions and transcription factor overexpression, lipids from *C. pacifica* were extracted and neutral TAGs were converted into fatty acid methyl ester (FAME) biodiesel and analyzed using Gas Chromatography-Mass Spectrometry (GC-MS). Following lipid extraction and esterification, the components of each final biodiesel were identified, and the FAMEs were sorted based on their chain length and degree of unsaturation. All final samples contained trace amounts of “non-biodiesel” nonpolar lipid components such as cholesteryl esters. However, purity remained high across all samples, ranging from 85-97% without any added purification steps (**Figure 7**, **Supplementary Figure 6**). Samples grown under nitrogen-starvation conditions exhibited a reduced proportion of “non-biodiesel” components in the final samples compared to those grown in normal media with percent purity of biodiesel FAME components ranging from 95-98% (**Figure 7**, **Supplementary Figure 6**). This observation aligns with existing reports on common microalgal responses to nitrogen-deprivation; growth of microalgae in nitrogen-deplete media has been observed to induce neutral lipid TAG production due to potential shifts in cellular biosynthetic pathways that normally rely on the presence of nitrogen^49–51^. Though the increased TAG accumulation in nitrogen-starved samples correlated with higher overall FAME biodiesel purity, the high purity of biodiesel from all samples with overexpressed transcription factors is a promising indicator of their potential for biodiesel production. A major challenge in use of wildtype microalgae as feedstocks for biodiesel is the lack of TAG accumulation compared to other plant-based oil feedstocks^52^. This was observed in previous experiments to produce biodiesel from wildtype evolved *C. pacifica*, where fatty acids needed to initially be isolated from total lipid extracts to produce biodiesel at this level of purity^30^. The high purity of biodiesel combined with the observed increased TAG accumulation under different conditions in this experiment suggests that these newly modified strains of *C. pacifica* could be a promising source of microalgal TAGs for biodiesel production.

**Figure 7:**
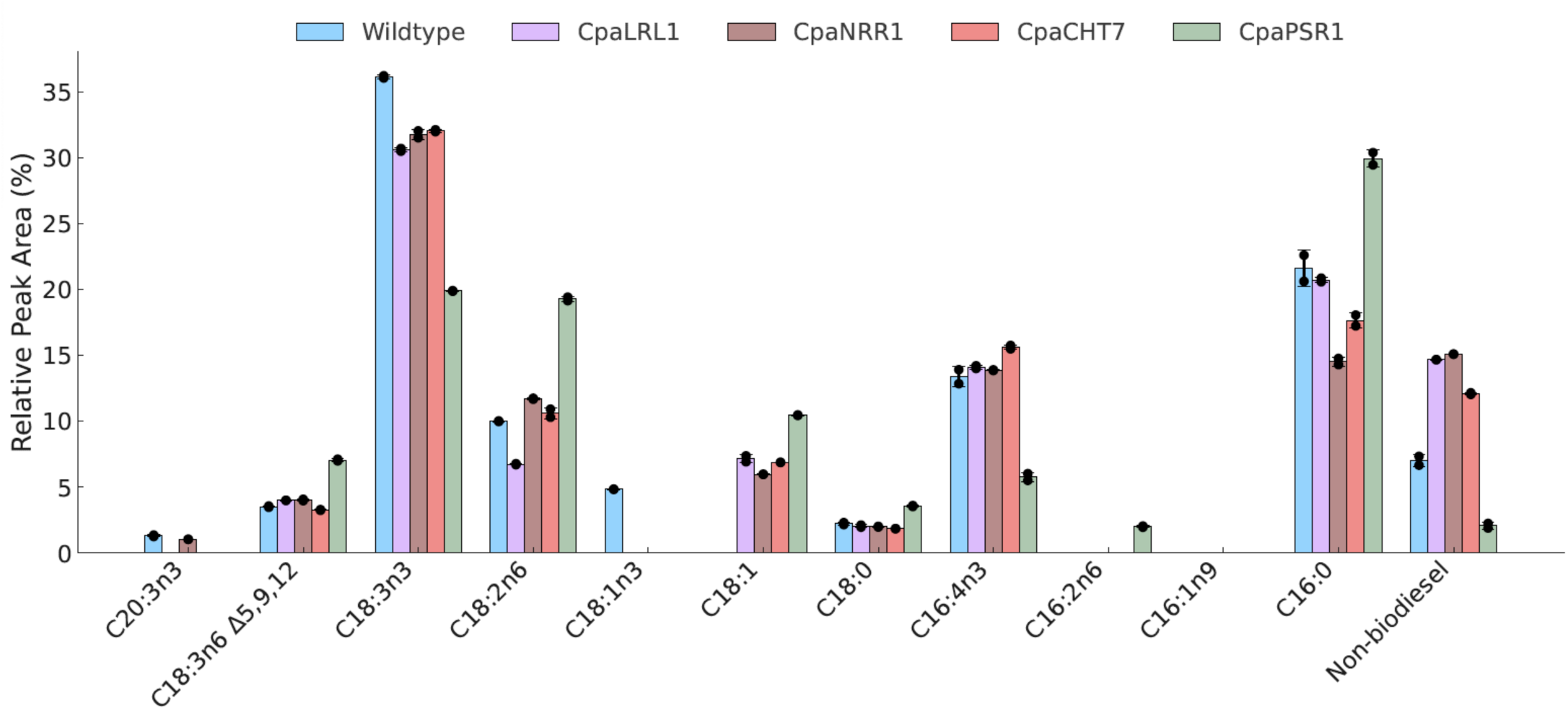
GC-MS analysis of algae-derived FAME biodiesel. Comparison of biodiesel composition of overexpressed lines with evolved wildtype *C. pacifica* in normal minimal media through relative peak area integration percentages in gas chromatography-mass spectrometry analysis (n = 2).

For initial analysis, the FAME profile of the original evolved *C. pacifica* strain developed by Gupta & Molino et al. (CC-6190 402wt x 403wt) was compared to CpaLRL1, CpaNRR1, and CpaCHT7 grown in minimal media (**Figure 7**). While no significant differences in fatty acid profiles were seen between these samples, a secondary comparison with biodiesel from CpaLRL1, CpaNRR1, and CpaCHT7 grown under nitrogen-deplete media revealed promising changes in the fatty acid composition (**Supplementary Figure 6**). Notably, a decrease in the production of C18:3n3 and C16:4n3 polyunsaturated biodiesel in favor of the production of C16:0 and C18:1 biodiesel was seen. The biodiesel composition of the original evolved *C. pacifica* strain was also compared to FAME profiles from CpaPSR1 grown under normal media conditions (**Figure 7**). A similar reduction in polyunsaturated fatty acids was seen in the CpaPSR1 compared to the evolved strain, and the C16:0 FAME production in the CpaPSR1 grown in normal media was the highest across all samples (**Figure 7**).

## Discussion

Transcription factor-based metabolic engineering has proven to be highly effective in enhancing biomass quality in microalgae^19^. Notably, numerous studies have employed both exogenous and endogenous transcription factors to successfully increase lipid production across various microalgal strain^36,43,47,53^. In this study, we have demonstrated a high-throughput pipeline for identifying and validating lipid-enhancing transcription factors in a new microalga. We used *C. pacifica* as an example species due to its great potential as a commercially relevant green extremophile alga. Previously, we showed that overexpression of an exogenous soybean-based *Dof* transcription factor led to increase in lipid accumulation in *C. pacifica* under nitrogen-deprived conditions compared to the wild-type strain^30^. Here, we identified more potent endogenous transcription factors that result in higher lipid accumulation in minimal media compared to the wild type. This discovery not only avoids cell starvation which can lead to lower biomass productivity, but also saves time and resources needed to create nitrogen-deprived conditions for large-scale cultivation.

This study successfully presented an in-silico high-throughput pipeline designed to identify lipid-enhancing transcription factors in microalgae (**Figure 2**). Our approach involved predicting all transcription factors within the genome of the target species and subsequently comparing these predictions with a curated database of validated transcription factors from ten different genera, as reported in the literature^21,34,43,54–59^. The novelty of this approach lies in its ability to streamline the identification process, significantly reducing the time and resources typically required for experimental validation. By applying and validating this pipeline in the newly discovered species *C. pacifica*, we demonstrated its robustness and reliability. The successful identification and experimental validation of lipid-enhancing transcription factors in *C. pacifica* not only showcase the efficacy of our approach but also instill confidence in its application to other commercially relevant microalgae species. Given the increasing demand for sustainable biofuels and bioproducts, this pipeline could play a pivotal role in advancing microalgal biotechnology by accelerating the discovery of key genetic regulators that enhance lipid production.

Analysis of transcription factor distributions across species (**Figure 1**) supports existing data that multicellular land plants possess a significantly larger TF repertoire compared to unicellular algae like *Chlamydomonas reinhardtii*^60^. The clear demarcation between algae and higher plants in the PCA plot underscores the evolutionary divergence in the TF complexity and possibly reflects the distinct regulatory needs of unicellular and multicellular organisms. In contrast to other green algae, *C. pacifica’s* alignment with *C. reinhardtii* underscores potential evolutionary conservation in their TF composition. This relationship implies that the breadth of prior work involving *C. reinhardtii* can inform studies on *C. pacifica*, aiding green algae biotechnological advancements. Conversely, *C. pacifica* exhibited the presence of TFs from the Trihelix family, a group not as extensively studied in green microalgae. In higher plants, Trihelix TFs are known to coordinate genes related to salicylic acid metabolism, a pathway integral to plant stress responses^61,62^. Identifying Trihelix TFs in *C. pacifica* opens new avenues for research, particularly in understanding their functional roles in microalgae. This finding could potentially bridge gaps in knowledge regarding the regulatory mechanisms in green algae, especially those linked to environmental stress responses.

Next, to evaluate the full potential of the predicted TFs, we overexpressed them in *C. pacifica* using the β2 tubulin promoter sequence with hygromycin selection. We successfully obtained transformants for four out of the nine transcription factors that were cloned for overexpression. The partial success might be due to the β2 tubulin promoter being too strong, resulting in excessive expression of the transcription factor and potential lethality for cell growth^63^. As *C. pacifica* is a newly discovered species with limited molecular tools, this study did not prioritize testing alternative promoters to explore potential variation among different transformation clones. However, the identified TFs are promising candidates for further testing under inducible promoters^64,65^. Nonetheless, we did not observe any negative impact on the transgenic lines of the four tested TFs (**Figure 4**). Using cytometry and confocal microscopy, we observed improved or unchanged lipid accumulation in all four-transcription factor overexpressing lines under normal or nitrogen-deprived media (**Figure 5, 6**). To further boost lipid accumulation, future efforts can employ mutagenesis and breeding techniques on these transgenic lines^66^.

Previously, the LRL1 mutant was known to suppress growth and lipid accumulation under phosphorus stress due to its role in lipid remodeling^58^. In our study, we demonstrated that overexpression of CpaLRL1 does not affect cell growth and leads to higher lipid accumulation under nitrogen-deprived conditions, indicating its role in managing nitrogen stress. Similarly, the NRR1 mutant in *C. reinhardtii* was shown to reduce TAG levels by 50% under nitrogen starvation^59^. In accordance, our study found that overexpressing this transcription factor increases lipid accumulation under nitrogen-starvation conditions. Moreover, the CHT7 mutant is known to impact lipid degradation and cell regrowth when nitrogen is resupplied^57,67^. In our study, we observed that overexpression of CpaCHT7 increases lipid production under normal media conditions. Additionally, the PSR1 knockdown line has been shown to inhibit lipid accumulation and play a role in starch metabolism. Overexpression of PSR1 in a cell-wall-deficient strain increases starch content without changing or reducing neutral lipid levels^43^. This also aligns with findings from another study showing that PSR1 overexpression in normal wild-type strains of *C. reinhardtii* also results in higher intracellular lipid content^44^. We found that overexpressing PSR1 in *C. pacifica* increases lipid accumulation when nutrients were not limiting, but did not improve lipid accumulation when nitrogen is limited. Collectively, these results suggest different roles for PSR1 transcription in cell-wall-deficient and normal wild-type strains regarding lipid biosynthesis.

To fully demonstrate the potential of the engineered lines, we proceeded to use their biomass for biodiesel production. A current challenge in producing high-quality diesel from biobased resources lies in feedstocks’ natural fatty acid composition compared to the ideal chemical composition of high-performance diesel^68^. An ideal biodiesel formulation should have a high cetane number (CN), indicating a faster rate of fuel ignition upon injection into the engine and complete combustion while maintaining high oxidative stability and moderate viscosity^69^. Per the regulations set in ASTM standard D6751-24 for bio-based middle distillate fuels (jet fuel and diesel fuel), biodiesel fuel blend stocks should possess a minimum cetane value of 45, kinematic viscosity between 1.9-6 mm^2^/s at 40°C, and a minimum oxidative stability of 3 hours at 110°C^70^. The FAME analysis of the evolved *C. pacifica* in this study is consistent with our previous findings and closely resembles the profile of *C. reinhardtii*, with the highest compositions being C16:0 and C18:3n3^30,71^. Moreover, the observed shifts in fatty acid composition in samples with overexpressed lipid-associated transcription factors are relevant towards improved properties of biodiesel. Comparing above-mentioned metrics to the fatty acids produced by *C. pacifica* in this study, the increased production of C16:0 and C18:1 in the CpaPSR1 samples and nitrogen-starved CpaLRL1, CpaNRR1, and CpaCHT7 have favorable implications for the production of high-quality algae biodiesel. The metrics for C16:0 methyl palmitate (CN=86, 4.38 mm^2^/s, >24h stability) and C18:1 methyl oleate (CN=57, 4.51 mm^2^/s, 2.79h stability) demonstrate improvements towards the ASTM standard for biofuel compared to the C18:3n3 methyl linolenate (CN=23, 3.14 mm^2^/s, 0.00h stability) that comprised the majority of the biodiesel from the initial evolved *C. pacifica* strain^69,72^. Paired with the high lipid accumulation observed under normal media conditions, the high abundance of C16:0 methyl ester biodiesel from the CpaPSR1 sample indicates that this strain of *C. pacifica* could be a useful tool for industrial-scale production of sustainable fuel. However, the finding differs from the fatty acid composition observed in PSR1 overexpression lines in *C. reinhardtii*, which could be attributed to species-specific differences or variability among clones^44,73,74^.

Nonetheless, leveraging these prior findings of the functionality of PSR1, CHT7, NRR1, and LRL1 in *C. reinhardtii* in the context of new genomic resources for *C. pacifica* reveals how these transcription factors can be used in tandem to influence the metabolic processes of a new algal strain. Complete pathways for TAG and starch biosynthesis were found in the *C. pacifica* genome based on homology to other *Chlamydomonas* genomes and KEGG orthologs (**Figure 8**). *C. pacifica* genes relevant to TAG biosynthesis are identified in **Supplementary Table S6**. PSR1 and CHT7 appear to improve starch and TAG accumulation, respectively, by downregulating genes associated with their utilization^43^. Our confocal microscopy supports prior observations that CHT7 improves lipid accumulation even in the presence of a nitrogen source (**Figure 6**). Prior transcriptomic work on NRR1, LRL1, and PSR1 suggests that these transcription factors drive TAG accumulation primarily through the upregulation of the Kennedy pathway rather than the upstream production of free fatty acids^43,58,59^. Thus, additional transcription factors driving type II fatty acid synthase systems could be used with the transcription factors in this study to improve TAG biosynthesis further. CHT7 preventing utilization of TAGs may explain why it appears to be more effective in nutrient rich conditions (**Figure 5a**) compared to the other cell lines, as additional TAGs produced from upregulation by PSR1, NRR1, or LRL1 might cause additional TAG hydrolysis and lessen TAG buildup. Consistency of transcription factor behavior between *Chlamydomonas* species is encouraging for industrial exploitation, as novel strains with valuable characteristics such as light and halotolerance can undergo genetic engineering using the wealth of knowledge gained from *Chlamydomonas reinhardtii*.

**Figure 8:**
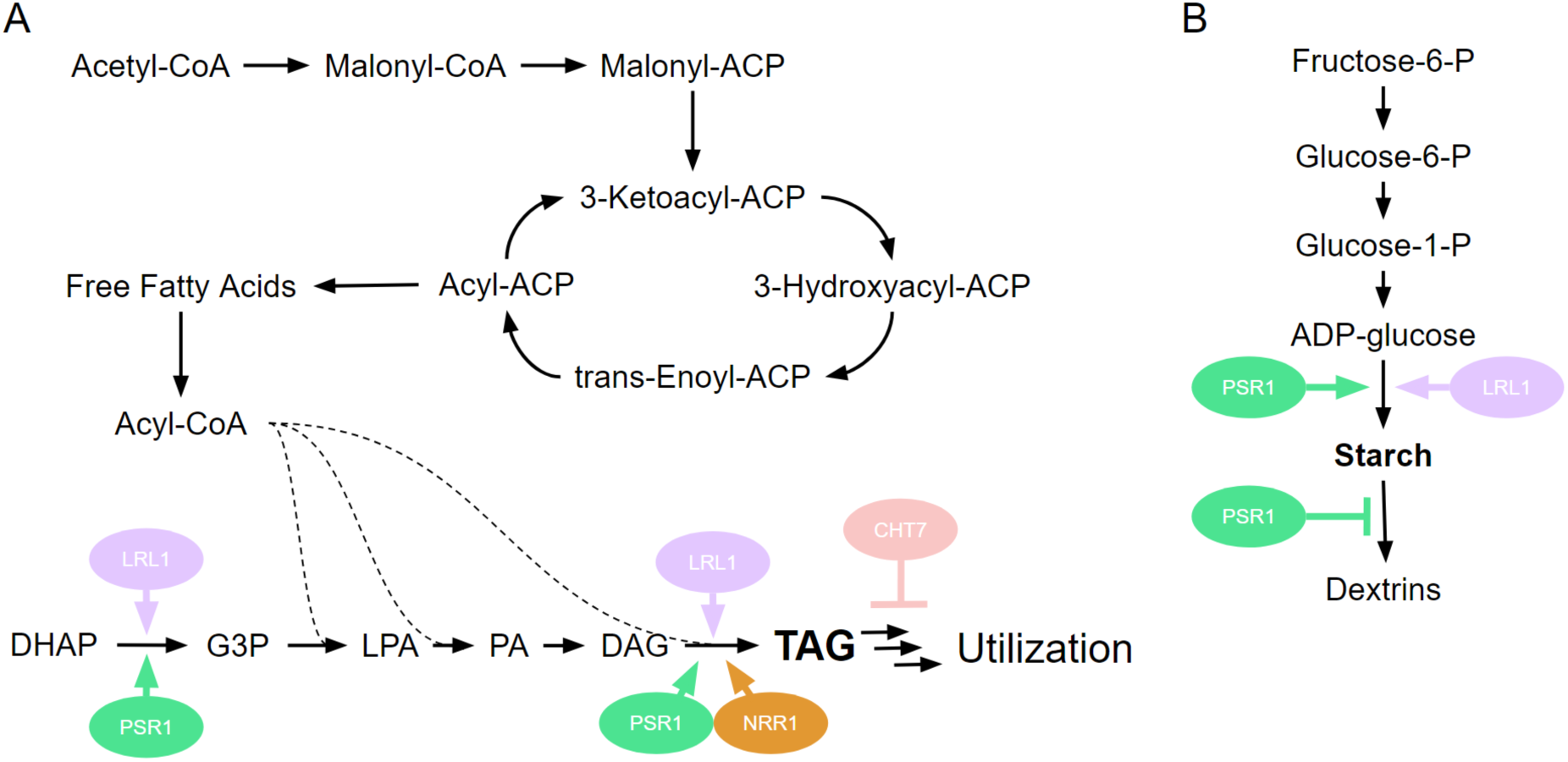
Proposed model showing how transcription factors shape triglyceride and starch accumulation in *Chlamydomonas pacifica* through increased production and decreased degradation. The metabolic pathways for (**A**) TAG and (**B**) starch biosynthesis and initial degradation in *C. pacifica*. Arrows from TFs represents putative upregulation, and perpendicular tipped lines represent putative downregulation. Abbreviations: ACP, acyl carrier protein; DHAP, dihydroxyacetone phosphate; G3P, glycerol-3-phosphate; LPA, lysophosphatidic acid; PA, phosphatidic acid; DAG, diacylglycerol; TAG, triacylglyceride.

In this study, we have tested and demonstrated several transcription factors that enhance lipid production within the same species. Future studies could focus on generating additional lines for these transcription factors and performing comprehensive lipidomic analyses to assess whether they can be leveraged to synthesize specific fatty acid lipid chains. If specificity is found, transcription factor-based tuning of different fatty acid chains could have significant applications for industries other than biofuels, including food, cosmetic, and nutraceutical.^7,75–77^

## Conclusion

This study underscores the potential of utilizing endogenous transcription factors to enhance lipid accumulation in *Chlamydomonas pacifica*, positioning it as a prime candidate for large-scale biodiesel and biomaterial production. By identifying and overexpressing key transcription factors—CpaLRL1, CpaNRR1, CpaCHT7, and CpaPSR1—we achieved substantial increases in lipid biosynthesis compared to the wild-type strain, with differential roles observed under nitrogen-deprived conditions. The significant increase in lipid accumulation in CpaPSR1 overexpressing strains highlights its commercial relevance, and the successful conversion of these lipids into biodiesel demonstrates their practical applicability. This approach not only advances our understanding of lipid biosynthesis in extremophile algae but also offers a robust framework for transcription factor-focused metabolic engineering in other commercially important microalgae species, contributing to sustainable bioenergy solutions and a circular bioeconomy. Future studies could explore the impact of combinatorial overexpression of the tested transcription factor on lipid production and other commercially valuable metabolites.

## Methods

### Algae strain and genomic data

The wild-type strain of *Chlamydomonas pacifica* has been submitted to the Chlamydomonas Resource Center under the identifiers CC-5697 and CC-5699^23^. The datasets employed in our research, encompassing genome assembly, protein sequences, coding DNA sequences for all proteins, and predicted transcript data, are available at the following URL: https://doi.org/10.5281/zenodo.10811182 or were sourced from Molino et al.^29^. For all experiments, we utilized the evolved *C. pacifica* strain developed by Gupta & Molino et al. (CC-6190 402wt x 403wt)^30^. We cultivated the cultures following the conditions outlined by Gupta & Molino et al. Specifically, we used High-Salt Media with acetate (HA) and urea as the nitrogen source at a pH of 10.5. The culture media recipe was adapted from the work of Molino et al.^29^ The cells were grown under continuous 24-hour light at approximately 25°C, with a photon flux of 125 µE/m²/s, on shaken tables rotating at 125 RPM. Biomass for lipidomics and biodiesel analysis was obtained via settling and centrifugation after algal growth in 10L media carboys. Prior to lipid extraction, biomass samples of *C. pacifica* were flash frozen in a dry ice/isopropanol bath for lyophilization to remove excess aqueous media (LabConco FreeZone 2.5, 0.6 mbar reduced 498 pressure, -52°C).

### Prediction of transcription factors

Our methodology for predicting transcription factors involved the use of three key bioinformatics tools: PlantTFDB (version 5), TAPscan, and iTAK^33,78,79^. These tools were selected to leverage their specific strengths in TF identification and classification. A notable distinction among these databases is the quantity of Chlorophyte genomes each contains. Specifically, PlantTFDB (version 5) and TAPscan (version 3) include 16 and 13 Chlorophyte genomes, respectively, in contrast to iTAK, which houses just six algae genomes. Our lipids and starch TFs predictions were cross-referenced with a list compiled by Gupta et al., to ensure comprehensiveness and accuracy^21^. For multiple sequence alignment, we utilized EMBL-EBI Clustal Omega, chosen for its efficiency and reliability in aligning protein sequences^80,81^. The alignments facilitated a comparative analysis of the predicted TFs^82^. Subsequently, phylogenetic trees were generated using the Simple Phylogeny tool, providing insight into the evolutionary relationships of these TFs^80,82^. 10 *C. pacifica* transcription factors’ closest sister node was previously experimentally validated starch or lipid, and these TFs were selected for further experimental validation. General functionalities for transcription factors were annotated based on categories from the March 2022 version of the Database of Clusters of Orthologous Genes (COGs) predicted using eggNOG-mapper (version 2)^83,84^. Bar plots for transcription factor classes and COG categories were visualized using R version 4.2.1 and R package ggplot2 (version 3.4.4.)^85^.

### Principal component analysis and network plot

Transcription factor distributions for algal species visualized via PCA (**Figure 1C**) were sourced from Hu *et al.*^32^ and PlantTFDB (version 5)^33^. The PCA was generated using the sklearn package in Python (version 3.10.9), with additional preprocessing using the “StandardScaler” function to normalize TF abundance between species. A sequence similarity network of TFs from 16 chlorophyte genomes in PlantTFDB^33^ (**Figure 1E**) was constructed using all-vs-all blastp with a minimum percent identity cutoff of 50%^86^. Transcription factors that did not reach this similarity cutoff to another TF were not pictured. The resulting network was visualized in R version 4.2.3 using the graphopt layout of igraph v. layout^87^.

### Conserved domain analysis

Our study’s conserved domain analysis entailed comparing predicted transcription factors with established ones from the literature. We utilized the protein sequences of *Arabidopsis* WRINKLED1 (AtWRI1) and sunflower WRINKLED1 (HaWRI1) as presented in Sánchez et al.^48^. Additionally, we referenced conserved domain information for PHOSPHORUS STARVATION RESPONSE 1 (PSR1) TF in *C. reinhardtii* and *A, thaliana* from Rubio et al.^88^. The multiple sequence alignments were visualized using the pymsaviz (version 0.4.2) package in Python, available at https://pypi.org/project/pyMSAviz/. This approach enabled the efficient alignment and visualization of similarities and conserved domains between our predicted TFs and those from *Arabidopsis* and sunflower.

In our analysis, we performed a multiple sequence alignment of transcription factors from *C. pacifica* and related species, focusing on the key functional regions (**Supplementary Figure 4a, b**). In the case of CrePSR or AtPHR1, the key function region corresponds to the alignment position of 247-300 and 447-489 based on *A. thaliana*’s PHR1. Whereas in the case of AtWRI1 and HaWRI1, it corresponds to alignment positions of 107-177 and 212-271. We concatenated these regions to emphasize conserved domains and calculated percentage identity from the concatenated sequences. We used permutation testing with numerous iterations (n = 100,000) to determine statistical significance, randomly shuffling amino acids within the regions of interest across all sequences. This generated a distribution of alignment scores under the null hypothesis, from which we derived a p-value, indicating the probability of the observed alignment occurring by chance.

### Identification of lipid and starch metabolism gene orthologs and promoter motif analysis

In our analysis, we focused on identifying and validating the orthologs of genes involved in starch and triacylglyceride metabolism in *Chlamydomonas reinhardtii*, as characterized by Bajhaiya et al. and Shin et al.^43,89^. Utilizing the gene sequences from this study, we conducted local NCBI BLASTP searches to find corresponding orthologs in *C. pacifica* using Biopython (version 1.81) package in Python with a cutoff e-value = 0.001^90^. The functional annotation of the identified orthologs was conducted using the EggNOG tool, with the e-value and score as calculated by EggNOG^91^. Further annotation was performed using the KEGG Automatic Annotation Server to confirm that *C. pacifica* contains a complete pathway for starch and sucrose metabolism (pathway: ko00500) and fatty acid biosynthesis (pathway: ko00061)^92,93^. For promoter sequences, the analysis focused on the 1000 base pairs upstream of the transcription start site. During this promoter analysis, we searched for motifs that were similar to the consensus motif of the PSR1 transcription factor, as identified in the PlantTFDB database (TF ID: Cre12.g495100.t1.2)^33^. Transcription factor binding site analyses for PSR1 and CHT7 used PlantTFDB-provided transcription factor binding motifs, scanned against the *C. pacifica* genome using FIMO version 5.5.5^94^.

### Vector design, algae transformation, and Reverse Transcription-Quantitative Polymerase Chain (RT-qPCR) Reaction

The vector assembly utilized genetic components from the genome of *C. pacifica*. The β-Tubulin A 2 gene promoter was employed to regulate the expression of hygromycin (used as a selection marker) and the endogenous transcription factor genes. All the vectors (pJPCHx1_CpaDpWRI1, pJPCHx1_CpaAtWRI1, pJPCHx1_CpaMYB6, pJPCHx1_CpabZIP1, pJPCHx1_CpaSPL12, pJPCHx1_CpaPSR1, pJPCHx1_CpaCHT7, pJPCHx1_CpaNRR1, pJPCHx1_CpaLRL1) have been submitted to the Chlamydomonas Resource Center at the University of Minnesota in St. Paul, Minnesota and the plasmid sequences can be accessed in the following URL: https://doi.org/10.5281/zenodo.13287080.

For the algal transformation, plasmid DNA was first digested with KpnI and XbaI enzymes (New England Biolabs, Ipswich, MA, USA). Subsequently, the DNA fragments were purified using the Wizard SV Gel and PCR Clean-up System (Promega Corporation, Madison, WI, USA) without fragment separation. The DNA concentration was then determined using the Qubit dsDNA High Sensitivity Kit (Thermo Fisher Scientific, Waltham, MA, USA), and the DNA amount for each transformation was adjusted to ensure 1 µg of payload DNA, based on the payload-to-backbone DNA ratio. The transformation was carried out via electroporation following the protocol by Molino et al. (2018) ^95^. Transformed cells were selected on HA agar plates containing 30 µg/mL hygromycin.

To confirm successful transformation, Colony-PCR was conducted using primers from Gupta & Molino et al., targeting a 277 base pair segment of the hygromycin gene (forward primer: TGATTCCTACGCGAGCCTGC; reverse primer: AACAGCTTGATCACCGGGCC) (**Supplementary** Figure 5)^30^. The housekeeping gene primers target the ATP2 gene, amplifying an 86 bp segment using the following sequences: forward primer – ACGGTGGTTTCTCTGTGTTC, and reverse primer – CACACCCGACTCAATCATCTC. The results were then verified through sequencing.

To evaluate the abundance of transcription factor transcripts, total RNA was extracted from the samples using the Direct-zol RNA Miniprep Plus Kit (Zymo Research). Complementary DNA (cDNA) was subsequently synthesized using the AzuraFlex cDNA Synthesis Kit (Azura Genomics) with a combination of random hexamers and oligo(dT) primers, ensuring broad coverage of the RNA transcriptome. Quantitative PCR (qPCR) was performed using the SsoAdvanced Universal SYBR Green Supermix (Bio-Rad) on a Bio-Rad thermocycler. The qPCR protocol included an initial denaturation and polymerase activation step at 98 °C for 3 minutes, followed by 40 amplification cycles of 15 seconds at 95 °C for denaturation and 1 minute at 60 °C for annealing and extension. A melt curve analysis was conducted from 65 °C to 98 °C, increasing by 0.5 °C every 5 seconds, to confirm the specificity of the amplified products. ATP2 served as the housekeeping gene for normalization, and the relative expression levels of target genes were calculated using the 2^−ΔΔCT^ method^96^. The primers used for amplification were: ATP2 (housekeeping gene)–Forward: 5′-ACGGTGGTTTCTCTGTGTTC-3′, Reverse: 5′-CACACCCGACTCAATCATCTC-3′; CpaLRL1-Forward: 5′-GAGCGGTTCAATCACCATTTG-3′, Reverse: 5′-CGGTTGCCGAAACGTAGA-3′; CpaNRR1-Forward: 5′-TGCAACATGGACCACGAA-3′, Reverse: 5′-GCATGTGAACGCCTTACAAC-3′; CpaCHT7-Forward: 5′-CTTCCAAGCAACGACAACAC-3′, Reverse: 5′-GGGAATTCCACCTGGAAAGA-3′; CpaPSR1-Forward: 5′-CAGATGGACCTGCAGAAGAAG-3′, Reverse: 5′-CATGAGGCTGGCGATGTAG-3′.

### Lipid staining and flow cytometry

We followed the method outlined by Gupta and Molino et al.^30^. In brief, a 50 µL sample of the culture was combined with 110 µL of buffer to make a final volume of 160 µL. Next, 1.6 µL of 1 mM Nile red in DMSO was added to this mixture. Three replicates were used for each transgenic line (or wildtype) per nitrogen condition. The cells were incubated at 40°C for 10 minutes using a Tecan Infinite M200 Pro plate reader (Tecan Group Ltd., Männedorf, Zurich, Switzerland). After incubation, the samples were analyzed with a Beckman Coulter CytoFLEX Cytometer. Intracellular lipids were detected by measuring Nile Red fluorescence using the PE channel (585/42 bandpass filter) with an excitation/emission of 551/636 nm and a gain setting of 1. For each sample, 10,000 events were recorded at a flow rate of 30 µL/min. The visualized RFU threshold was selected to be one standard deviation above average wild-type nitrogen-replete RFU (approx. 7.7×10⁴), surpassed by 3.32% of control cells. A Wilcoxon ranked-sum test with p < 0.05 was used to determine increased lipid production in transgenic lines compared to wild-type controls.

### Confocal Microscopy

The sample slides were prepared using Frame-Seal™ Slide Chambers filled with agar enhanced with media. Cells were positioned on the agar and covered with a cover slip to seal them. The confocal microscopy was conducted at the UCSD School of Medicine Microscopy Core using a Leica STED microscope. Chlorophyll was detected with an excitation wavelength of 405 nm and an emission range of 630-680 nm. For Nile red, the excitation wavelength was 470 nm, and the emission range was 475-575 nm. An ANOVA with p < 0.05 was used to determine if cell growth or chlorophyll levels were significantly different between cell lines.

### Lipidomics analysis

Lipids analysis were conducted at UCSD Lipidomics Core facility. Lipids were extracted following protocols described by Murawska et al., Hartler et al. and Quehenberger et al.^97–100^. 8-10mg of algae biomass was homogenized into 200uL 10% methanol. A mix of deuterated lipid standards (Equisplash, Avanti) was added to 150uL biomass homogenate and extracted via BUME. Briefly, samples were extracted using 250 μL of butanol/methanol (BuOH/MeOH), 250 μL of heptane/acylacetate, and 250 μL of 1% acetic acid. The lipid-containing top layer was collected, solvents were removed, and the lipids were reconstituted in 50 μL of buffer (18:1:1 IPA/DCM/MeOH). Chromatographic separation was performed using a Thermo Vanquish UPLC system with a Cortecs T3 (C18) column (2.1 mm x 150 mm, 1.8 μm particle size), using a binary solvent system. Five microliters (about 7.5% of total biomass) of each sample were injected. Lipid identification and quantification were conducted using a Thermo Q Exactive mass spectrometer operating in data-dependent MS/MS acquisition mode. LipidSearch and LDA were used for data analysis. The relative abundance of TAG metabolites was analyzed through co-injection with an EquiSPLASH LipidoMIX 15:0-18:1-d7-15:0 TG quantitative mass spec internal standard. The relative abundance is calculated by analyte intensity/internal standard intensity.

### Biodiesel

Algal biodiesel was synthesized from total lipid extracts using acidic Fischer esterification to produce FAMEs. The masses of dried samples were recorded and two technical replicates per transcription factor and growth condition were used for subsequent lipid extraction and biodiesel synthesis. For each replicate, 0.25g of dry biomass was resuspended into 1mL of deionized water and the total lipids were extracted using the Bligh-Dyer extraction protocol^101,102^. Following lipid extraction into chloroform, the solvent was evaporated under air before proceeding with biodiesel synthesis. Each dried lipid sample was treated with an excess of 1M hydrochloric acid in methanol with stirring at 65°C for 2h. Biodiesel was recovered from this mixture via liquid-liquid extraction into hexane. The solvent was removed under ambient evaporation and the final liquid biodiesel was diluted to 2mg/mL in hexane for GC-MS characterization. GC-MS analysis of biodiesel was performed on an Agilent 7890A GC system connected to an Agilent 520 5977C GC/MSD. Samples were separated on an Agilent HP-5MS UI 30m x 0.25mm x 0.25um GCMS column with hydrogen as carrier gas and a gradient of 3°C /min from 70°C to 250°C over 60 mins. The mass-to-charge ratio and NIST23 library were used to identify peaks associated with methyl ester biodiesels. Integration values from associated peaks were used to determine the abundance of different biodiesel FAMEs within each sample. The plots were generated using GraphPad Prism.

## Statistics and Reproducibility

Comprehensive details of the statistical analysis are provided in the respective section, with significance determined at p-values less than 0.05. To ensure reproducibility, the experimental design has been thoroughly described in the Methods section, and data has been made available wherever feasible.

## Data Availability

We have thoroughly included links to the deposited data throughout the Methods section.

## Ethics declarations

### Authors Contributions

**AG**: conceptualized the research framework, designed the computational methodologies, developed the software tools, performed the data curation, executed the computational analyses, and contributed to writing, editing, coordinating, and conceptualization of the original draft. **AO**: developed the software tools, executed the computational analyses, and contributed to the writing and editing of the original manuscript. **JVDM**: conceptualized the research framework, contributed to experiments and writing and editing of the original manuscript. **KWF**: contributed to experiments and writing and editing of the original manuscript. **MT**, **KK, EES**: contributed to experiments. **YT-T**: contributed to reviewing the original manuscript. **MDB**, **SM**: contributed to the writing and editing of the original manuscript and secured financial support for the research.

## Acknowledgments

The authors would like to extend their sincere thanks to Prof. Stefan Rensing (University of Freiburg) and Romy Petroll (Max Planck Institute for Biology Tübingen) for their assistance in conducting the TAPscan prediction on our dataset. The authors express their gratitude to Jennifer Santini and Marcy Erb from the UCSD School of Medicine Microscopy Core, supported by the NINDS P30NS047101 grant, for their assistance with confocal microscopy. The authors extend their gratitude to Dr. Oswald Quehenberger, Aaron Armando, and Milda Simonaitis from the UCSD Lipidomics Core for their invaluable assistance with lipidomics analysis. Additionally, the authors are grateful for the support provided by the United States Department of Energy, the California Center for Algae Biotechnology, and the University of California San Diego.

