## Supplementary Figures for "Identification and Overexpression of Endogenous Transcription Factors to Enhance Lipid Accumulation in the Commercially Relevant Species *Chlamydomonas pacifica*"

### Supplementary Data

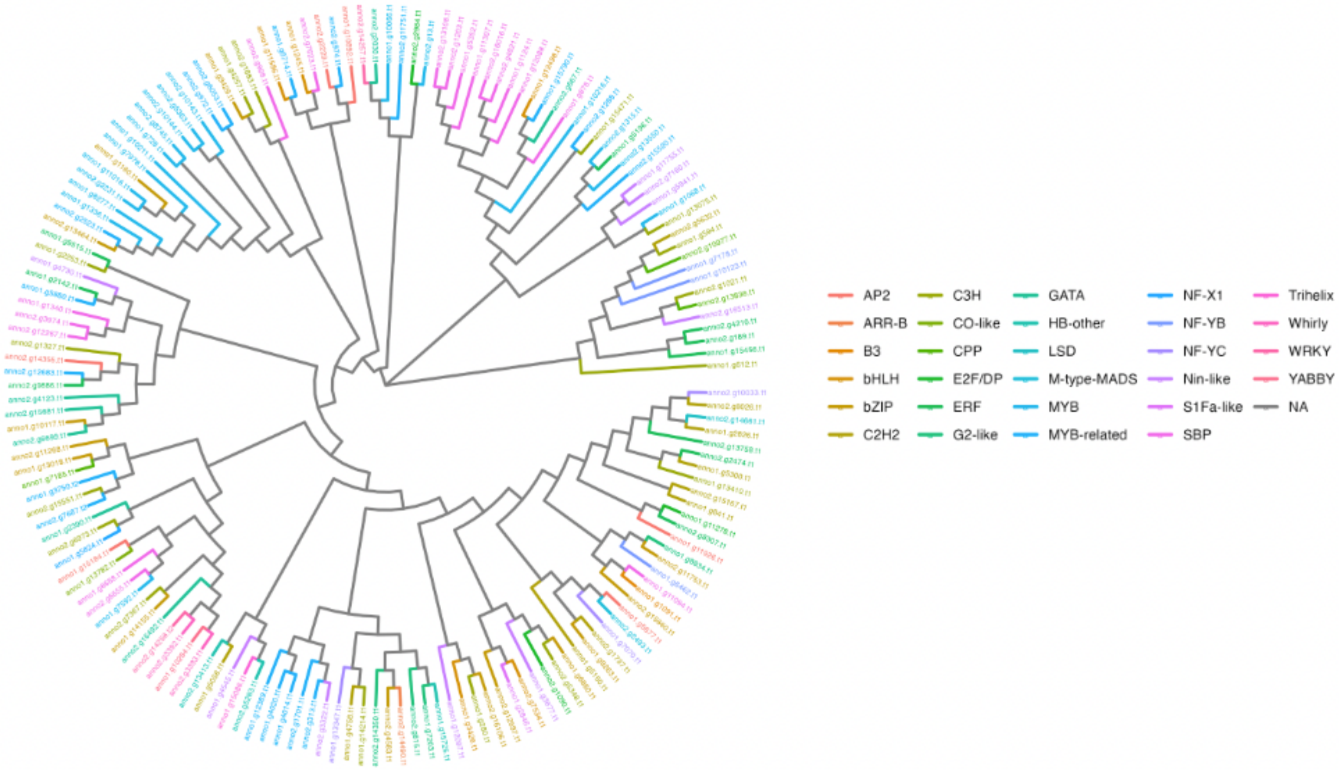


**Supplementary Figure 1**: Circular phylogenetic tree depicting the evolutionary relationships of transcription factors in *Chlamydomonas pacifica*.

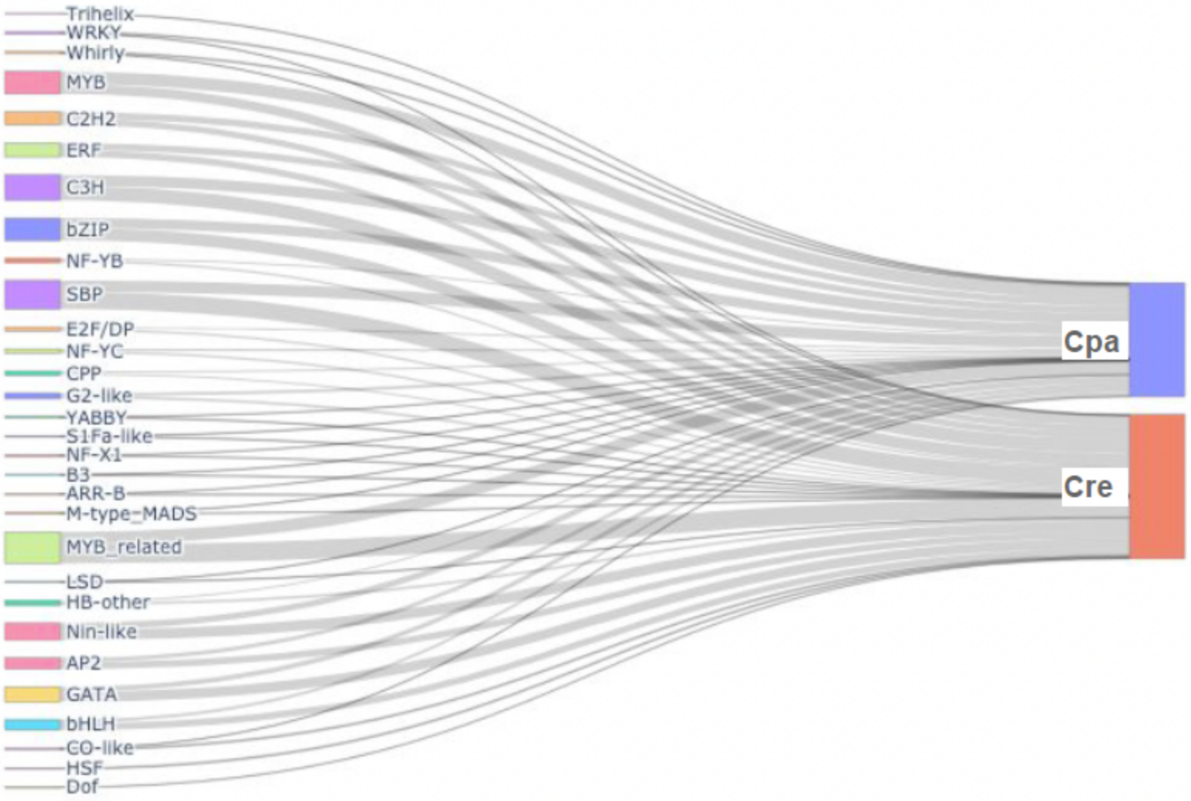


**Supplementary Figure 2**: A Sankey plot depicting the transcription factor profile comparison between *Chlamydomonas pacifica* (Cpa) and *Chlamydomonas reinhardtii* (Cre) for different transcription factor families. The notations for transcription factor families were sourced from PlantTFDB (v5.0).

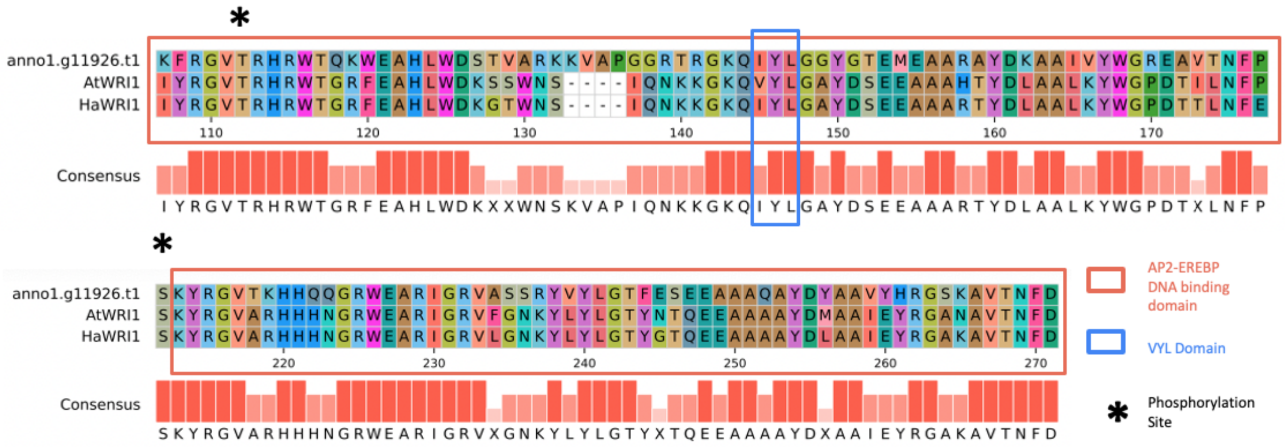


**Supplementary Figure 3: Multiple sequence alignment showing conserved domains in WRI1.** Alignment showing WRI1 transcription factor from *A. thaliana* (AtWRI1), *Helianthus annuus* (HaWRI1)), and a predicted homolog in *Chlamydomonas pacifica* (anno1.g11926.t1), showcasing the conserved AP2 DNA-binding domains, VYL transcriptional activation motif, and two phosphorylation sites

**Supplementary Figure 4a**: Multiple Sequence Alignment (MSA) fasta file for AtPHR1, CrePSR1, and anno1.8834.t1.

```
>AtPHR1
MEAR------PVHRSGSRDLTRTSSIPSTQKPSPVEDSFMRSDNNSQLMSRPLGQTYHLL
SSSNGGAVGHICSSSSSGFATNLHYSTMVSHEKQQHYTGSSSNNAVQTPSNNDSAWCHDS
LPGGFLDFHETN------PAIQNNCQIEDGGIAAAFDDIQKRSDWHEW-----ADHLITD
D-----DPLMSTNWNDLLLETNSNSDSKDQKTLQIPQPQIVQQQPSPSVELRPVSTTSSN
SNNGTGKARMRWTPELHEAFVEAVNSLGGSERATPKGVLKIMKVEGLTIYHVKSHLQKYR
TARYRPEPSETGSPERK---------------LTPLEHIT--------------------
------------------------------------------------------------
---------------SLDLKGGIGITEALRLQMEVQKQLHEQLEIQRNLQLRIEEQGKYL
QMMFEKQNSGLTKGTASTSDSAAKSEQEDKKTADSKEVPEEETRKCEELESPQPKRPKID
N-----------------------------------------------------------
------------------------------------------------------------
------------------------------------------------------------
------------------------------------------------------------
-----------------------------------------------
>CrePSR1
MDKAERAAGGPNAASEDDWLLEFWPEPAADFPAPVAPMLSQHQDAAQLPE-AM-------
PQQQGLALG--------GYGL------------------------TQQ----PSDFMQTG
MP-GFDAFSSGKAATLGLPLLADPQRASTDGASALMNAAQQSSEYMLAPGMGGMPHLLAP
SVGTALPGTGHTGFADLSMGG----MAGGIP--GLGGPGI-MHG--------QYFMQPQR
AATGPAKSRLRWTPELHNRFVNAVNSLGGPDKATPKGILKLMGVDGLTIYHIKSHLQKYR
LNIRLPGESGLAGDSADGSDGERSDGEGGVRRATSLERADTMSGMAGGAAAALGRAGGTP
GGALISPGLAGGTSSTGGMAAGGGGGGGLVTEPSISRGTVLNAAGAVATAAPAAAAPAGG
SAAVKRPAGTSLSSGSTASATRRNLEEALLFQMELQKKLHEQLETQRQLQLSLEAHGRYI
ASLMEQEGL--TSRLPELSGGA-PAA-----------APVAAGGAAGGMIAPPPPQQQLQ
HQPQLLQPQGSLPAGGSSE-AHAAAGAGTMVVHQQQQQHVHHHHQQQQVQMQQHARHCDT
CGAGGAGGAPSGGSSMQQLQAAEQQRTELVVAGRLGSMPAPASSSPLAGQAHQQQPLAGG
AAHLVHVHSHTPGGQPHVQHQDAFAGAATAAAHASPGLPQSHSHLLPADLSSNAGPDTSA
GQIKPEPDMSQQQQQQEQQEAEQLAQGLLNDSSAGAGAVSGSDGGGLGDFDFGDFGDLDG
GAQGGLLGPGDLIGIAELEAAAAHEQQQEQEHDPLDADRAKRQRVEP
>anno1.g8834.t1
------------------------------------------------------------
------------------------------------------------------------
------------------------------------------------------------
-------------------MH----QPATFPSLGVLQPGLLPQG--------HFFAQPQR
P-AAPAKSRLRWTPELHNRFVSAVNQLGGPDKATPKGILKLMGVDGLTIYHIKSHLQKYR
LNIRLPGETIQGDSEPTVSNTTHGQG----------------------------------
------------------QGQGQGSKGG--------------------------------
-----VRRGVSASSGSTASATRRNLEEALLFQMDLQKKLHDQLETQRQLQLSLEAHGRYI
ASLMEQEGL--TRLPPELTSGGAPAG-----------ATPSAGRGPAGA--PQLGGAH--
-----------MTTGGRLEQQHLLV----------------------------NAGIC--
------------------------------------------------------------
------------------------------------------------------------
------------------------------------------------------------
-----------------------------------------------
```

**Supplementary Figure 4b**: MSA fasta file for AtWRI1, HaWRI1, and anno1.g11926.t1

```
>anno1.g11926.t1
MDTDYMQPNTGGGHNPWGLDTNAFFSSLGGVSGAGTRRDR-SMQDPPNSAFDASTLFHNP
HRMSGDEGEAGAGGTTRRNSSASGPLSVALAPGTTPVVRSKNGGTSKFRGVTRHRWTQKW
EAHLWDSTVARKKVAPGGRTRGKQIYLGGYGTEMEAARAYDKAAIVYWGREAVTNFPMAE
YEDIMEELQGLNKETVVSMLRRGSCGFSRGASKYRGVTKHHQQGRWEARIGRVASSRYVY
LGTFESEEAAAQAYDYAAVYHRGSKAVTNFDVRNYIDMVTGDAGILSGA---APNAAVAY
HVS--M------------------------------------------------------
------------------------------------------------------------
------------------------------------------------------------
----
>AtWRI1
```

```
---------------------------------MKKRLTTSTCSSSPSSSVSSSTTTSSP
IQS---EAPRPKRAKRAKKSSPSGDK------SHNPTSPASTRRSSIYRGVTRHRWTGRF
EAHLWDKSSWNS----IQNKKGKQVYLGAYDSEEAAAHTYDLAALKYWGPDTILNFPAET
YTKELEEMQRVTKEEYLASLRRQSSGFSRGVSKYRGVARHHHNGRWEARIGRVFGNKYLY
LGTYNTQEEAAAAYDMAAIEYRGANAVTNFDISNYIDRLKKKGVFPFPVNQANHQEGILV
EAKQEVETREAKEEPREEVKQQYVEEPPQEEEEKEEEKAEQQEAEIVGYSEEAAVVNCCI
DSSTIMEMDRCGDNNELAWNFCMMDTGFSPFLTDQNLANENPIEYPELFNELAFEDNIDF
MFDDGKHECLNLENLDCCVVGRESPPSSSSPLSCLSTDSASSTTTTTTSVSCNYLFQGLF
VGSE
>HaWRI1
---------------------------------MKRRRLSPTSSSSCSSS------SSCI
NDH---QIPKSKRPCSRPHNKNTTQI------SNQNQNAETTRRSSIYRGVTRHRWTGRF
EAHLWDKGTWNS----IQNKKGKQIYLGAYDSEEAAARTYDLAALKYWGPDTTLNFEIDT
YKKDIEEMEKLSKDEYLASLRRRSSGFSRGVSKYRGVARHHHNGRWEARIGRVLGNKYLY
LGTYGTQEEAAAAYDLAAIEYRGAKAVTNFDISIYADRLKNLPQTQT-----TPESTEVV
ASP--------KHEQNEEVLHDINS---QQQQEQQQQQPQQQQEVALGEKEELMPGIHNL
D------F-PQTVEEEHPWSLCLDSS-YNLLPVPHFSFDKSGDQLLDLFDDTGFEDNIDF
MFEGLISCENEVEKVHACINSPSS----SSSLSSS---------SRTNSLSCPVSHY---
----
```

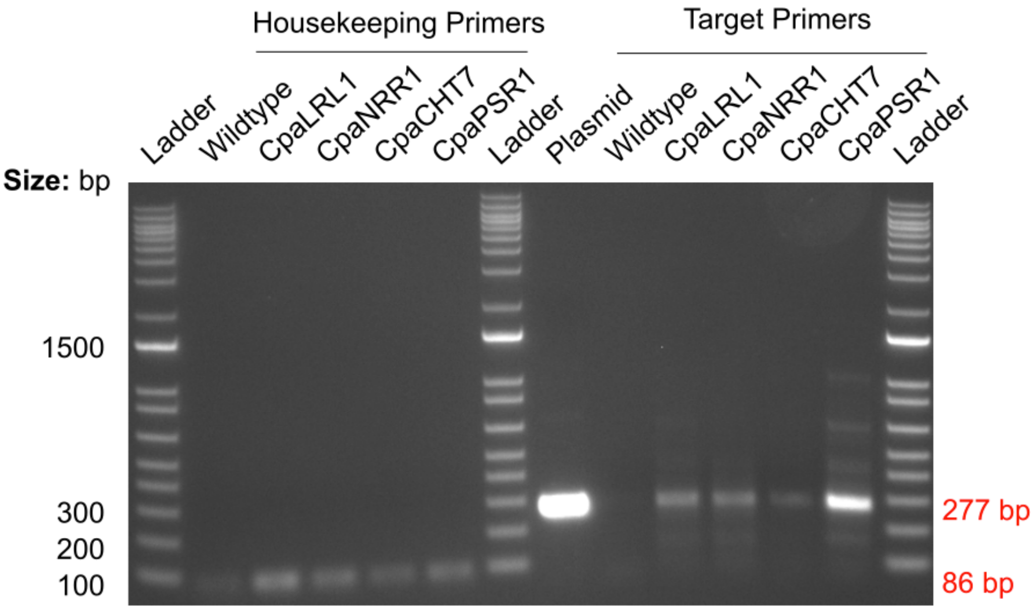


**Supplementary Figure 5: Colony PCR of transformants.** PCR was performed to detect the presence of the hygromycin resistance gene (target) and the ATP2 gene (housekeeping control). A plasmid containing the hygromycin gene was used as a positive control. A 1 kb Plus DNA ladder was used as a molecular size marker.

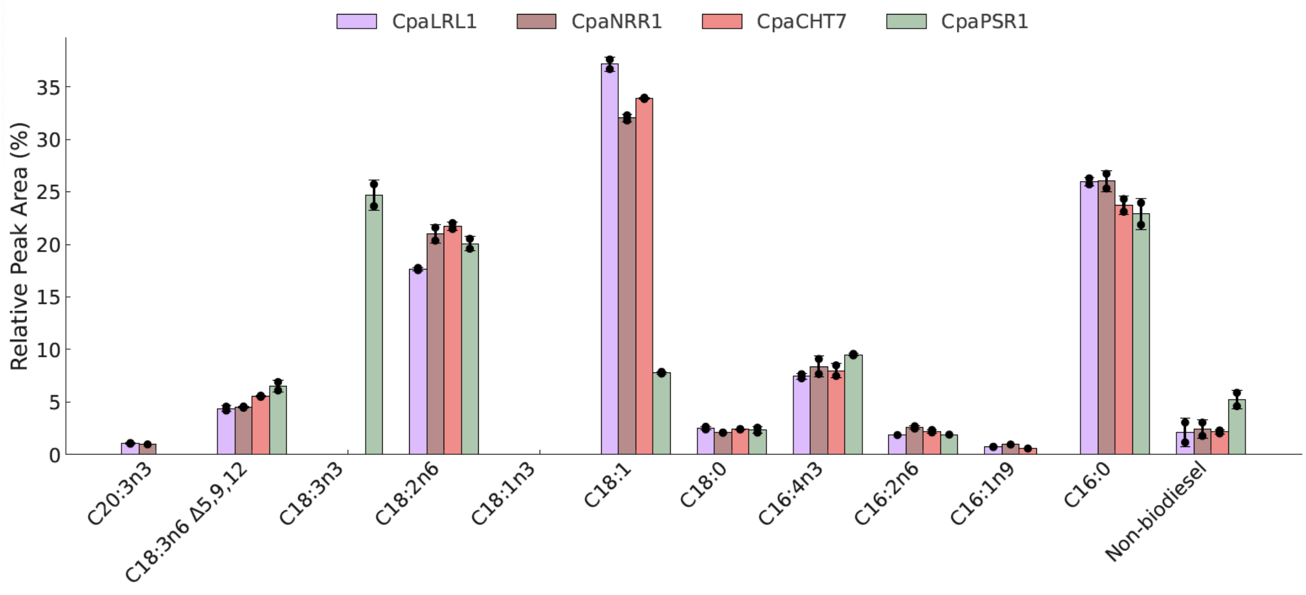


**Supplementary Figure 6: GC-MS analysis of algae-derived FAME biodiesel of overexpressed lines in nitrogen-deprived media**
